# Dependency of LKB1-inactivated lung cancer on aberrant CRTC-CREB activation

**DOI:** 10.1101/2021.01.09.425982

**Authors:** Xin Zhou, Jennifer W. Li, Zirong Chen, Wei Ni, Xuehui Li, Rongqiang Yang, Huangxuan Shen, Jian Liu, Francesco J. DeMayo, Jianrong Lu, Frederic J. Kaye, Lizi Wu

## Abstract

Lung cancer with loss-of-function of the *LKB1* tumor suppressor is a common aggressive subgroup with no effective therapies. *LKB1-*deficiency induces constitutive activation of cAMP/CREB-mediated transcription by a family of three CREB-regulated transcription coactivators (CRTC1-3). However, the significance and mechanism of CRTC activation in promoting the aggressive phenotype of LKB1-null cancer remain poorly characterized. Here we observed overlapping CRTC expression patterns and mild growth phenotypes of individual CRTC-knockouts in lung cancer, suggesting functional redundancy of CRTC1-3. We consequently designed a dominant-negative mutant (dnCRTC) to block all three CRTCs to bind and co-activate CREB. Expression of dnCRTC efficiently inhibited the aberrantly activated cAMP/CREB-mediated oncogenic transcriptional program induced by LKB1-deficiency, and specifically blocked the growth of LKB1-inactivated lung cancer. Collectively, this study provides direct proof for an essential role of the CRTC-CREB activation in promoting the malignant phenotypes of LKB1-null lung cancer and proposes the CRTC-CREB interaction interface as a novel therapeutic target.

## Introduction

Lung cancer is the leading cause of cancer deaths in both men and women in the United States and worldwide (1–3). Global cancer statistics estimated 1,761,007 deaths due to lung cancer in 2018, contributing to about 20% of all cancer deaths (2). In 2020, there are an estimated 228,820 newly diagnosed lung cancer cases and 135,720 lung cancer deaths in the United States alone(1). Non-small cell lung cancer (NSCLC) accounts for approximately 85% of lung cancer cases and includes the major subtypes: lung adenocarcinoma, squamous cell carcinoma and large cell carcinoma (4). While small-molecule inhibitors targeted at driver gain-of-function gene mutations have achieved improved clinical outcomes over conventional cytotoxic therapy, they are currently available for only a subset of patients with lung cancer harboring specific mutations such as EGFR and ALK mutations (5–7). Cancer immunotherapy has emerged as one of the newest treatment options for NSCLC; however, the recent use of immune checkpoint inhibitors, such as those blocking the PD-1/PD-L1 checkpoint pathway, offer durable tumor responses to only a small population of patients generally with high tumor PD-L1 expression and/or high tumor mutational burden (4, 8). Therefore, effective treatments for the majority of lung cancer patients remain lacking.

Comprehensive genomic profiling has revealed the genetic landscape of lung cancer (9–11), identifying inactivating somatic STK11/*LKB1* (designated *LKB1* here) gene mutations as a common event in NSCLC. Somatic *LKB1* mutations arise preferentially in lung adenocarcinoma where they have been detected in up to 30% of cases (9, 11–13). In addition to gene mutations, *LKB1* can be inactivated by epigenetic silencing, post-translational modifications, or alterations in its interacting proteins (14–16). Lung cancer with LKB1 deficiency exhibits resistance to chemotherapy, targeted therapeutics and especially to immune checkpoint inhibitors in preclinical models and/or human patients (17–22). Therefore, the absence of targeted therapies and the lack of benefits of immune checkpoint inhibitors for this common aggressive lung cancer subtype require an urgent search for new therapeutic strategies.

*LKB1* was first identified as the cancer susceptibility locus for familial Peutz-Jeghers syndrome (PJS), which is characterized by mucocutaneous pigmentation and gastrointestinal hamartoma with an increased cancer risk (23, 24). Somatic inactivation of *LKB1* has now been observed in a variety of human cancers besides lung cancer. Importantly, LKB1 loss has been shown to promote cancer progression and increase metastatic potential in the genetically engineered mouse models of lung cancer, melanoma, pancreatic cancer and endometrial cancer (25–28). Also, *LKB1* mutations are associated with the suppressive immune milieu of the lung tumor microenvironment (21, 29). Thus, *LKB1* is a bona fide tumor suppressor gene. A better understanding of the pathogenic downstream signaling induced by LKB1 inactivation will facilitate the identification of rational therapeutic approaches.

The *LKB1* gene encodes a serine/threonine kinase that is essential for the activation of 14 downstream AMPK family members, such as AMP-activated protein kinase (AMPK) and salt-inducible kinases (SIKs) (30, 31). Therefore, LKB1 regulates multiple signaling pathways through its various substrates and plays critical roles in regulating cell polarity, metabolism and growth (30, 31). Consequently, LKB1 inactivation has the potential to promote tumorigenesis by deregulating downstream cell signaling, such as the defective LKB1-AMPK-mediated energy stress response which has been the focus of many studies (31). However, unlike loss of LKB1, loss of AMPK was found to reduce the growth of murine oncogenic Kras G12D-driven lung cancer (32), indicating that AMPK does not mediate LKB1’s tumor suppression function. To identify key signaling pathway(s) impacted by LKB1 inactivation in lung cancer, we previously performed an unbiased global gene expression profiling analysis and discovered that multiple cAMP/CREB-regulated targets, such as LINC00473, INSL4, NR4A1-3 and PTGS2, were highly expressed in human LKB1-null lung cancer cell lines and primary tumors (33, 34). The induction of these cAMP/CREB-mediated targets was linked with aberrant hyper-activation of the CRTC (CREB-regulated transcription co-activator) family in the context of LKB1 deficiency (33, 34). In addition, we previously generated an LKB1-null gene signature from 53 lung cancer cell lines to screen the Broad Institute Connectivity Map (CMAP) drug response database and the top 17 compounds that positively correlated with the LKB1-null gene signature were all compounds directly associated with CRTC activation (35). The CRTC family consists of three members, CRTC1, 2 and 3, which play important roles in metabolism, aging and cancer (36–39). These three CRTC proteins function as latent transcriptional co-activators and are normally sequestered in the cytoplasm. In response to cAMP and/or calcium signals, the family of three salt-inducible kinases (SIK1, 2, 3) are inactivated and/or phosphatases become activated, leading to CRTC dephosphorylation. Dephosphorylated CRTCs subsequently translocate to the nucleus and interact with the transcription factor CREB, activating CRE (cAMP-responsive element)-containing promoters. Since SIKs are dependent on LKB1 for its kinase activity, LKB1 deficiency impairs SIKs to phosphorylate CRTCs and consequently leads to an elevated level of unphosphorylated CRTCs, resulting in CRTC nuclear translocation and activation of CREB-mediated transcription. Therefore, the aberrant activation of the SIK-CRTC-CREB signaling axis may serve as a core driver event that underlies the aggressive phenotypes of LKB1-inactivated lung malignancies. This notion is further supported by recent CRISPR/Cas9-mediated gene editing studies revealing that knockouts of SIK1 and SIK3, but not of other AMPK family members, increased tumor growth in a mouse model of oncogenic KRAS-driven lung adenocarcinoma (40, 41). Therefore, SIKs mediate the major tumor suppressive effects of LKB1 in NSCLC. Moreover, CRTC2 was reported to promote tumor growth in LKB1-deficient NSCLC (42). However, the relative contributions of the three CRTC co-activators were not yet defined. Importantly, the role of the aberrant CRTC-CREB activation in LKB1-inactivated lung cancer and its underlying molecular mechanisms remained to be characterized.

In this study, we evaluated the significance and mechanisms of CRTC co-activators in lung cancers using CRISPR/Cas9-mediated knockouts of individual CRTCs and a pan-CRTC inhibitor that blocks all three CRTC co-activators’ ability to interact with the CREB transcription factor. Our *in vitro* and *in vivo* data provide direct evidence that CRTC activation plays an essential role in the growth of LKB1-deficient lung cancer cells and revealed that targeting gain-of-function CREB activation by interfering with the CRTC-CREB interaction is a potential effective strategy in treating LKB1-inactivated lung cancers.

## Results

### Three CRTC co-activators were expressed at varying levels in lung cancer cells

In our previous studies, we showed that human LKB1-null lung cancer cell lines and primary tumors exhibited aberrantly high expression of multiple cAMP/CREB targets, such as LINC00473, INSL4, NR4A1-3, and PTGS2 (33–35). The induction of these CREB-mediated transcriptional targets was due to LKB1 loss-induced activation of the CRTC co-activator family members, CRTC1, 2 and 3. A recent study reported CRTC2 as a critical factor in the growth of LKB1-deficient NSCLC (42). However, the individual functional contributions of the three CRTC family members and the general role of aberrant CRTC-CREB activation in lung malignancies remained undefined. Therefore, we first evaluated the expression patterns of the three CRTC co-activator family members by quantitative RT-PCR and Western blot analyses of SV40-transformed, non-tumorigenic human bronchial epithelial cells (BEAS-2B), 7 LKB1-wt and 6 LKB1-null NSCLC cell lines. We observed that the three CRTC genes were all expressed at varying levels, with CRTC2 and CRTC3 having higher relative expression than CRTC1 at the transcript level **(Figure 1A)**. Also, there were variable CRTC protein levels in all the cell lines examined and protein levels did not tightly correlate with their RNA transcript levels **(Figure 1B)**, suggesting potential post-transcriptional regulation. CRTC1 and CRTC2 exhibited predominantly faster migrating bands in all 6 LKB1-null cancer cells, consistent with enrichment of dephosphorylated forms in the setting of LKB1 deficiency. However, the mobility of CRTC3 appeared relatively unchanged in 4/6 LKB1-null cancer cell lines, suggesting a distinct pattern of post-translational regulation. In summary, we detected variable levels of expression of all 3 CRTC genes in immortalized human lung epithelial cells and lung cancer cells, suggesting the potential for both functional redundancies and unique properties. Since all three CRTCs are capable of co-activating CREB-mediated transcription (37, 38), we would need to inactivate each gene individually or all 3 together to determine the contribution of the CRTC co-activation to the transcriptional program in lung cancer cells with LKB1 deficiency.

**Figure 1:**
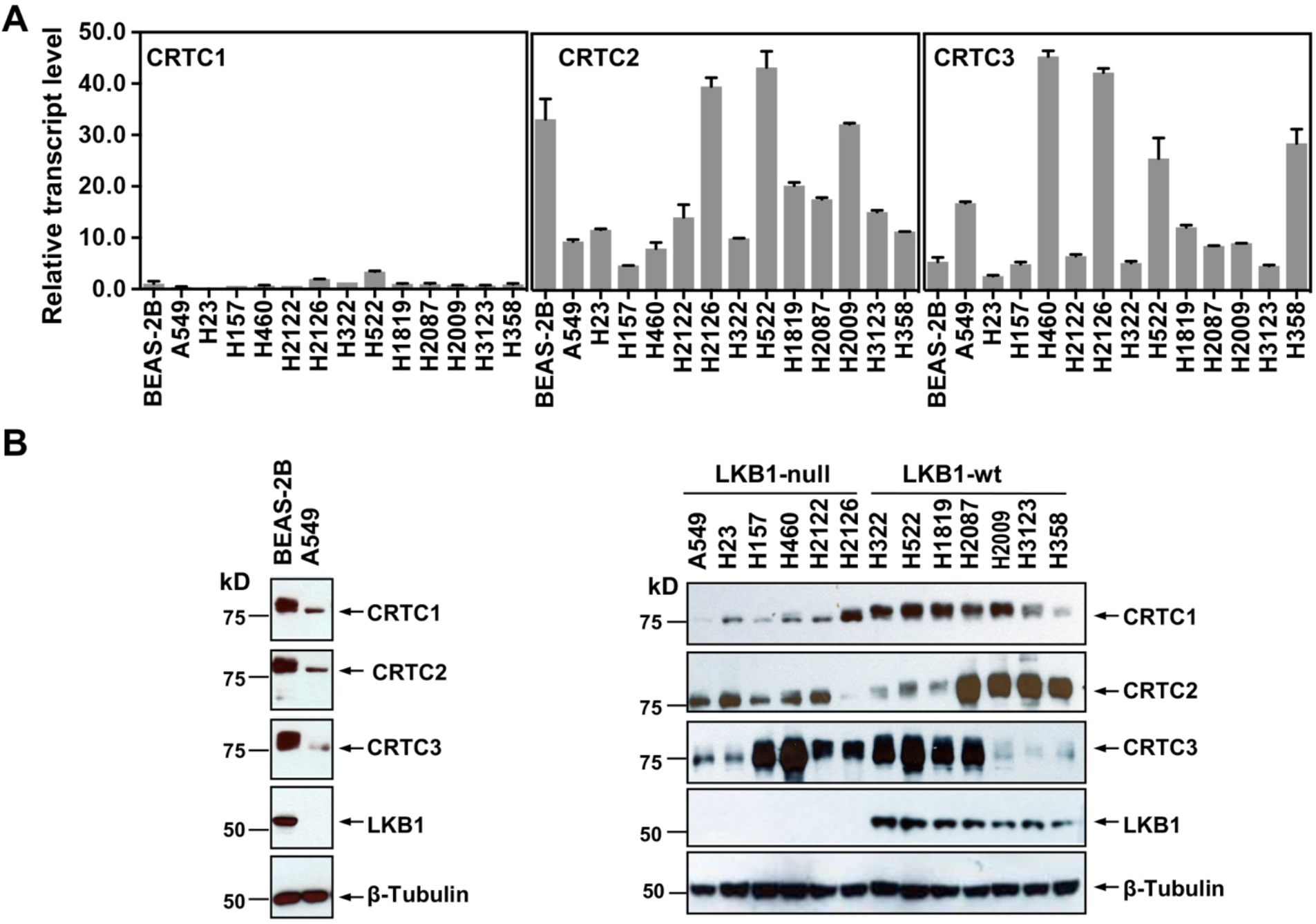
Three members of the CRTC co-activator family,*CRTC1*, *CRTC2* and *CRTC3*, are expressed at varying levels in human lung epithelial and cancer cell lines. **(A)** The transcript levels of three CRTC genes were determined by quantitative RT-PCR assays and normalized against the CRTC1 transcript level in BEAS-2B cells. **(B)** The protein levels of three CRTCs and LKB1 were detected by Western blotting. Blotting with anti-β-Tubulin was used as a loading control.

### CRISPR/Cas9-mediated knock-outs of individual CRTCs reduced expression of the CREB target genes and caused mild effects on NSCLC cell growth

To assess the importance of each CRTC family member in regulating lung cancer cell phenotype, we generated and characterized cells with individual CRTC knockouts. Specifically, human LKB1-null lung cancer A549 cells were transduced with lentiCRISPRv2 lentiviruses expressing single-guide RNAs (sgRNAs) for each CRTC gene or control sgRNA together with Cas9. Two independent, single knockout clones for each CRTC gene were then selected and CRISPR/Cas9-edited alleles with indels were further validated by genomic DNA sequencing **(Supplemental Figure 1).** We observed complete ablation of endogenous CRTC proteins in their respective knockout cells, as compared to the parental and control knockout cells by Western blotting **(Figure 2A)**. Upregulated CRTC1 protein levels were observed in response to CRTC2 knockout or CRTC3 knockout, indicating potential functional compensation. These individual CRTC knockout cells showed a reduction in expression of several CREB-mediated target genes, such as PDE4D, INSL4, LINC00473 and NR4A2, but not to the extent of their endogenous levels in LKB1-wt lung cancer H522 cells, as assayed by Western blotting or qRT-PCR assays **(Figure 2A, B)**. Individual CRTC knockout or control A549 cells were further assayed for cellular phenotypes, including cell viability, apoptosis, and anchorage-independent growth by trypan blue exclusion, annexin V/propidium iodide (PI) staining, and soft agar colony formation assays, respectively. We observed that knockout of each individual CRTC gene had only a mild effect on the numbers of viable cells, apoptotic cells, and colonies grown in soft agar (**Figure 2C, D, E**). These data indicate that the CRTC family members may be functionally redundant in regulating lung cancer cell proliferation, survival, and anchorage-independent growth.

**Figure 2:**
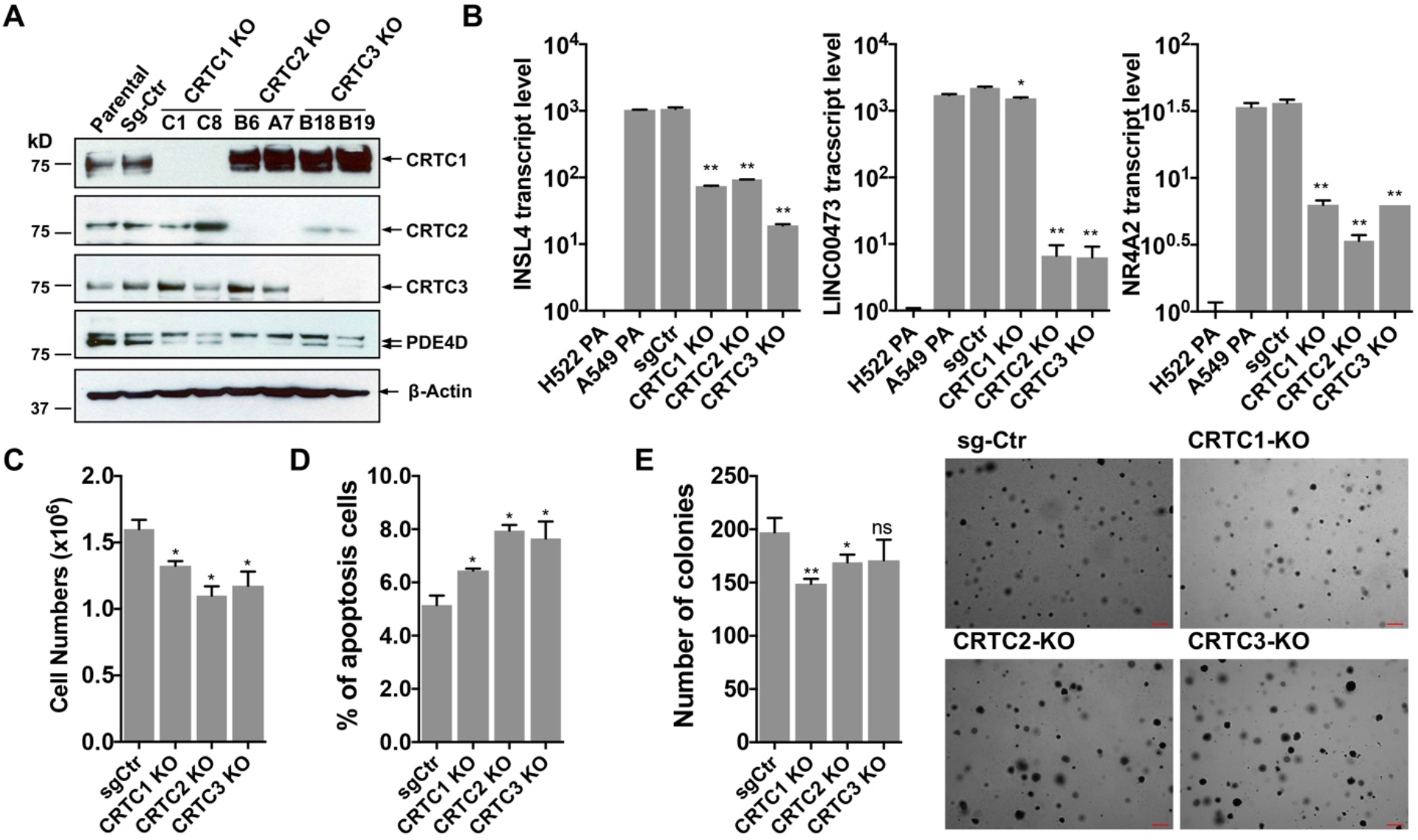
Individual knockouts of the CRTC family members in human LKB1-null lung cancer cells inhibit the CREB-mediated target gene expression and moderately affect cell viability and anchorage-independent growth. **(A)** Western blot analysis of endogenous CRTC proteins in parental A549 cells, A549 cells stably transduced with non-targeting sgRNA, and two independent single knockout clones for each CRTC1, CRTC2 or CRTC3. The protein level of a CREB target gene, PDE4D was also detected. Blotting with anti-β-ACTIN was used as a loading control. **(B)** The transcript levels of CREB-mediated target genes (INSL4, LINC00473 and NR4A2) were determined by RT-PCR assays (n=3). The LKB1-wt cells, H522 parental (PA) cells, were also analyzed. **(C,D)** Individual CRTC knockout or control cells were cultured at 3×10^5^ cells/well in the 6-cm plates for 96 hours. The viable cells were quantified by trypan blue exclusion assay (C), and the number of apoptotic cells was determined by staining with annexin V/propidium iodide (PI) followed by flow cytometry (D). **(E)** Control and CRTC knockout cells were cultured in soft agar for 14 days, and the resulting colonies were stained by crystal violet and photographed under microscope. The number of colonies was counted using ImageJ. Assays were performed in triplicate. One-way ANOVA test was used to calculate the p values (*p<0.05, **p<0.01, ns p>0.05).

### The dnCRTC (CRTC1 CBD-nls-GFP) functioned as a pan-inhibitor for the CRTC-CREB interaction and suppressed the CRTC-CREB signaling axis

Due to the potential functional redundancy of three CRTC coactivators in maintaining malignant cell behaviors of LKB1-null lung cancers, an approach of inhibiting all three CRTCs is required to assess the general role of aberrant CRTC activation in promoting tumorigenesis in LKB1-null lung cancer. The CRTC co-activators contain a highly conserved N terminal CREB binding domain (CBD) that is responsible for interacting with the transcription factor CREB, and a C terminal transcriptional activation domain (TAD) that is essential for transcriptional activation (36) **(Figure 3A)**. We, therefore, established a dominant negative approach of blocking the functions of all three CRTC co-activators by competing with endogenous CRTCs for CREB binding. Specifically, we generated a retroviral pMSCV-based dominant negative CRTC (dnCRTC) construct that expresses the CRTC1-CBD-nls-GFP chimeric protein, which contains the CBD of CRTC1 (1-55 aa) followed by a nuclear localization signal (nls, “PKKKRKV”) and EGFP. This CRTC1-CBD-nls-GFP protein was predicted to bind to CREB but lacks transcriptional activation, consequently interfering with the functions of endogenous CRTC co-activators through competitive CREB binding **(Figure 3A)**. We infected human LKB1-null lung cancer A549 cells with the CRTC1-CBD-nls-GFP or GFP (control) retroviruses and observed that CRTC1-CBD-nls-GFP was predominantly localized in the nuclear compartment, while the control GFP showed diffuse cytoplasmic and nuclear signals **(Figure 3B**). The CRTC1-CBD-nls-GFP chimeric protein showed an expected size of ~33 kDa **(Figure 3C)** and suppressed the ability of the three CRTC co-activators to activate the CREB-dependent transcription in cAMP response element (CRE)-containing promoter luciferase reporter assays **(Figure 3D).**Therefore, the CRTC1-CBD-nls-GFP chimeric protein functions as a dominant negative mutant for CRTC (dnCRTC), capable of blocking all three CRTCs to co-activate CREB-mediated transcription.

**Figure 3:**
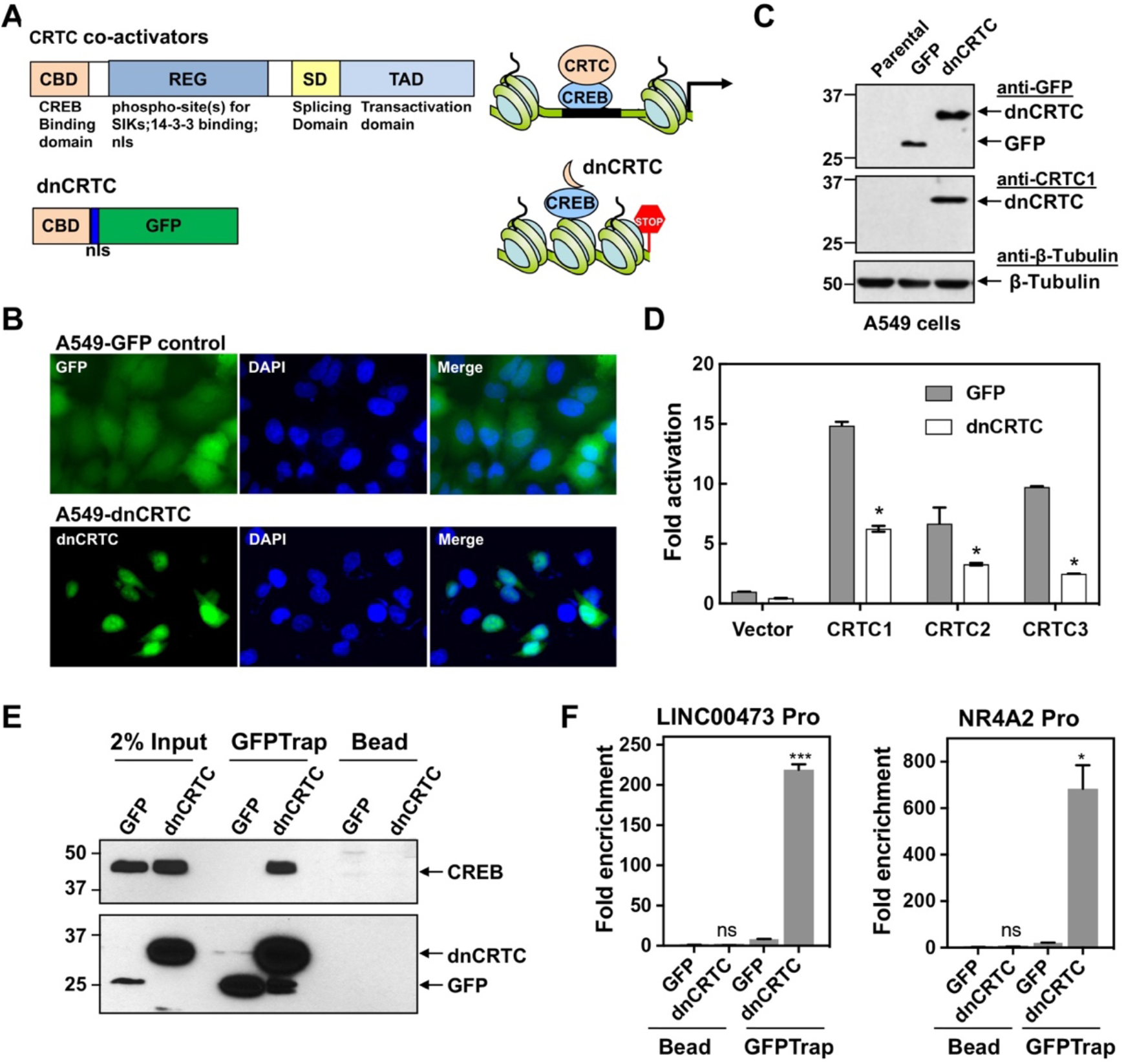
A dominant negative CRTC mutant (dnCRTC) interacted with CREB on the target gene promoters and blocked CRTC co-activation of CREB transcription. **(A)** A diagram of CRTC co-activator and dnCRTC was shown. The dnCRTC consists of CRTC1 (1-55aa) followed by a nuclear localization signal (nls) and GFP, cloned into the retroviral pMSCV vector. **(B)** A549 cells transduced with pMSCV-dnCRTC retroviruses showed that dnCRTC was predominantly localized in the nuclear compartment (lower), while A549 control cells transduced with pMSCV-GFP retroviruses showed both cytoplasmic and nuclear GFP signals (upper). DAPI stained for the nuclei. **(C)** Western blotting validated the expression of dnCRTC in transduced A549 cells. **(D)** Expression of dnCRTC blocked the abilities of CRTC1-3 to activate the pCRE-luc reporter in 293T cells (n=3). **(E)** dnCRTC interacts with CREB in the chromatin complex. Cells (GFP-expressing control and dnCRTC-GFP expressing cells) were crosslinked and chromatins were sonicated. GFP-Trap_A (anti-GFP VHH nano body coupled to agarose beads) were used for immunoprecipitation of dnCRTC-GFP proteins which were then blotted with anti-CREB and anti-GFP antibodies. Uncoupled agarose beads were used as control. **(F)** dnCRTC was enriched on the CRE regions of the promoters (n=3). Two-tailed student’s t-test was used to calculate the p values (*p<0.05, **p<0.01, ns p>0.05).

We next determined whether this dnCRTC interacts with the transcription factor CREB on endogenous CRE-containing gene promoters. Using chromatins prepared from dnCRTC- and control GFP-expressing A549 cells after cross-linking, we performed chromatin immunoprecipitation (ChIP) assay of dnCRTC or GFP using GFP-trap that consists of anti-GFP V_H_H nanobodies coupled to agarose beads (ChromoTek) or uncoupled agarose beads as negative control. Western blotting detected CREB in the dnCRTC-ChIP complex, but not in the control GFP ChIP complex **(Figure 3E)**, demonstrating a physical association of dnCRTC and CREB. We also observed that the DNA sequences spanning the CRE regions within the promoters of *LINC00473* and *NR4A2*, two genes known to be upregulated by CRTC-CREB activation due to LKB1 deficiency, were significantly enriched in the dnCRTC ChIP complex, but not in the control GFP ChIP complex by real-time PCR assays **(Figure 3F)**. Taken together, these data demonstrate that the dnCRTC (CRTC1-CBD-nls-GFP) mutant physically associates with CREB on the CRE-containing gene promoters and acts as a pan-inhibitor for all three CRTCs in co-activating CREB-mediated transcription.

### Inhibition of CRTC co-activators via dnCRTC effectively blocked the aberrant CREB-mediated transcriptional program in LKB1-null lung cancer cells

To evaluate the extent to which dnCRTC blocks the aberrant CRTC/CREB transcriptional program in LKB1-inactivated lung cancer, we profiled the transcriptomes of dnCRTC vs GFP-expressing cells to identify the affected downstream targets using an unbiased global screen. In brief, LKB1-null A549 lung cancer cells were transduced with dnCRTC and GFP retroviruses for 72 hours, and RNA was then isolated for gene expression profiling using Affymetrix GeneChip® Human Transcriptome Array 2.0. Two biological replicates were set up and expression of dnCRTC and control GFP was confirmed by Western blotting **(Figure 4A).** Using cut-off criteria of an absolute fold-change >= 2.0 and FDR p < 0.05, we identified a total of 274 dnCRTC-regulated differentially expressed genes (dnCRTC-DEGs), including 114 up-regulated and 160 down-regulated genes **(Supplemental Table 1)**; the heatmap and volcano plot were shown in **Figure 4B,C**. Since CRTCs are transcriptional co-activators, we next focused on the top downregulated dnCRTC-DEGs for the validation of the microarray results and confirmed that dnCRTC expression reduced the expression levels of multiple genes by qRT-PCR analysis **(Figure 4D**). These genes include known LKB1 target genes, such as *INSL4, CPS1, NR4A2, LINC00473, NR4A1, PTGS2, SIK1* and *PDE4D* and the overall high expression of these genes was associated with somatic mutations (SNPs and small INDELs) in *LKB1*, but not *KRAS* and *TP53*, in human TCGA lung cancers, particularly in adenocarcinoma (9, 10, 43) (**Supplemental Figure 2)**.

**Figure 4:**
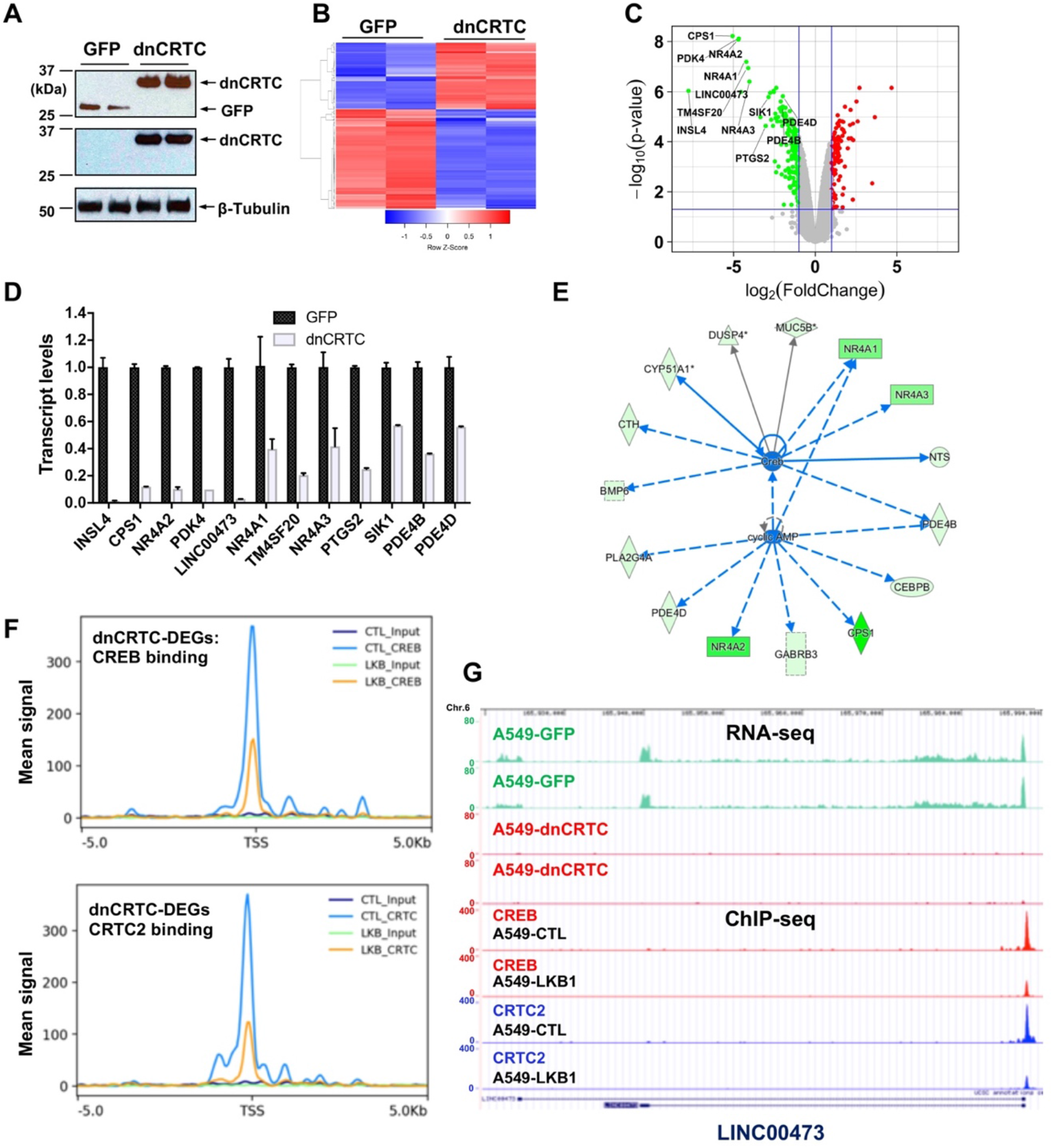
Gene expression profiling revealed dnCRTC repressed CRTC-CREB target gene expression. **(A)** Western blotting confirmed dnCRTC-expressing and GFP-expressing control cells. **(B, C)** The heatmap and volcano plots showed gene expression changes in dnCRTC-expressing and GFP-expressing cells. **(D)** The qRT-PCR analysis validated differential expressed genes (DEGs) in dnCRTC-expressing cells in A549 cells. **(E)** IPA analysis identified CREB and cAMP as upstream regulators for gene signature changes due to dnCRTC expression. **(F)** Analysis of CREB and CRTC2 binding of dnCRTC-DEGs in a ChIP-seq dataset. **(G)** CREB and CRTC2 binding peaks were shown in the LINC00473 target gene locus from the ChIP-seq analysis (lower panel). The mapped peaks of sequence reads from RNA-seq of A549-GFP and -dnCRTC cells was also shown (upper panel).

Ingenuity Pathway analysis revealed that CREB and cAMP are upstream regulators of gene expression changes observed in dnCRTC-expressing vs control A549 cells **(Figure 4E)**. Moreover, we analyzed the dnCRTC-DEGs for predicted CRE sites on their promoters (−3kb to 300bp from transcription start site) using the CREB Target Gene Database (44) and found that 169 of 274 (~61.7%) dnCRTC-DEGs contain predicted or experimentally verified CRE sites, which supports that dnCRTC affects a large set of CREB-regulated transcriptional loci (**Supplemental Table 1**). By incorporating a recently published ChIP-sequencing study (GSE128871) that investigated the genome-wide binding profiles of CREB and CRTC2 in LKB1-null A549 cells (42), we found that the dnCRTC-DEGs exhibited significant enrichment in CREB and CRTC2 binding around their transcription start sites (TSS), while the binding of CREB and CRTC2 was reduced upon reintroduction of *LKB1* (**Figure 4F**). This analysis revealed 97 of 274 (~35%) dnCRTC-DEGs (60 down-regulated and 37 up-regulated) having CREB-binding and CRTC2-binding peaks within −3 kb to 300bp from TSS; and 73 of 274 (~27%) dnCRTC-DEGs (45 down-regulated and 28 up-regulated) having both the CREB and CRTC2 binding peaks within - 500bp to 100bp from TSS (**Supplemental Table 1**). This list includes multiple known CRTC/CREB targets, such as *NR4A2, LINC00473* and *PTGS2.* A representative close-up view of the CREB and CRTC2 binding on the *LINC00473* gene locus was shown (**Figure 4G**). The mapped peaks of sequence reads from our RNA-seq re-analysis of A549-GFP and -dnCRTC cells were also shown. Overall, we identified a list of direct dnCRTC-regulated genes, which represent an extensive set of the potential critical mediators for CRTC-CREB activation in promoting lung cancer cell growth.

To gain further insights into the biological impact of dnCRTC expression, we performed gene set enrichment analysis (GSEA) of the transcriptomic data from dnCRTC-expressing vs control GFP-expressing A549 control cells using gene sets obtained from the Molecular Signatures Database. Several oncogenic gene signatures, including Shh-regulated gene set, RB loss/E2F1-regulated gene set, NFE2L2-regulated gene set, PDGF-regulated gene set, KRAS-regulated gene set, were found to be significantly altered with negative enrichment scores, indicating that a majority of genes in these oncogenic gene sets were significantly under-expressed in dnCRTC-expressing cells. Therefore, our dnCRTC mutant serves as a useful tool for blocking the extensive CRTC/CREB transcriptional program and oncogenic signaling. These data also suggest that dnCRTC expression has the potential to negatively impact the malignant behaviors of LKB1-deficient lung cancer cells.

### LKB1-null, but not LKB1-wt, NSCLC cells were sensitive to dnCRTC-induced inhibition of CRTC co-activators *in vitro*

To determine whether LKB1-null lung cancer cells depend on CRTC-CREB activation for growth and survival, we next assessed the functional impact of blocking the CRTC-CREB interaction via dnCRTC by analyzing the effect on lung cancer cell growth. We first performed competition assays using two LKB1-null (A549 and H157) and two LKB1-wt (H322 and H522) NSCLC cells. These cells were transduced with dnCRTC or control GFP retroviruses at an infection rate of ~40-60%, and then the percentages of GFP-positive cell populations were quantified at three-day intervals over a total of 24 days starting at day 3 following viral infection. We observed a progressively reduced percentage of LKB1-null cells (A549 and H157) that expressed dnCRTC, while the percent of the GFP-control cells remained stable **(Figure 5A)**. In contrast, the percentage of LKB1-positive cells (H322 and H522) was not significantly affected **(Figure 5A)**. These results showed that dnCRTC expression has a negative effect on the proliferation of LKB1-null tumor cells, but not of LKB1-positive cells, indicating that CRTC activation is critical for LKB1-null cell growth. We also sorted the GFP+ populations from dnCRTC- and GFP-transduced cells and performed functional comparisons. Expression of dnCRTC and GFP was first confirmed by Western blotting **(Figure 5B)**. We observed that dnCRTC expression induced a significant inhibition of cell growth in LKB1-null cells (A549 and H157), but not in LKB1-positive cells (H322 and H522) **(Figure 5C)**. Importantly, dnCRTC expression did not affect cell proliferation in normal lung epithelial cells (BEAS-2B) **(Supplemental Figure 3, A-C)**. Moreover, colony formation and soft agar colony formation assay showed that dnCRTC expression blocked colony-forming potential and anchorage-independent growth in LKB1-null cells (A549 and H157), but not in LKB1-positive cells (H322 and H522) **(Figure 5D,E)**. Expression of dnCRTC also had a similar negative effect on the growth of a mouse lung squamous carcinoma cell line, which was derived from a mouse model deficient of the tumor suppressors *LKB1* and *PTEN* (45) (**Supplemental Figure 3, D-G**). These results demonstrated that LKB1-null NSCLC cells are specifically sensitive to dnCRTC expression; therefore, they are highly dependent on the CRTC-CREB activation for growth.

**Figure 5:**
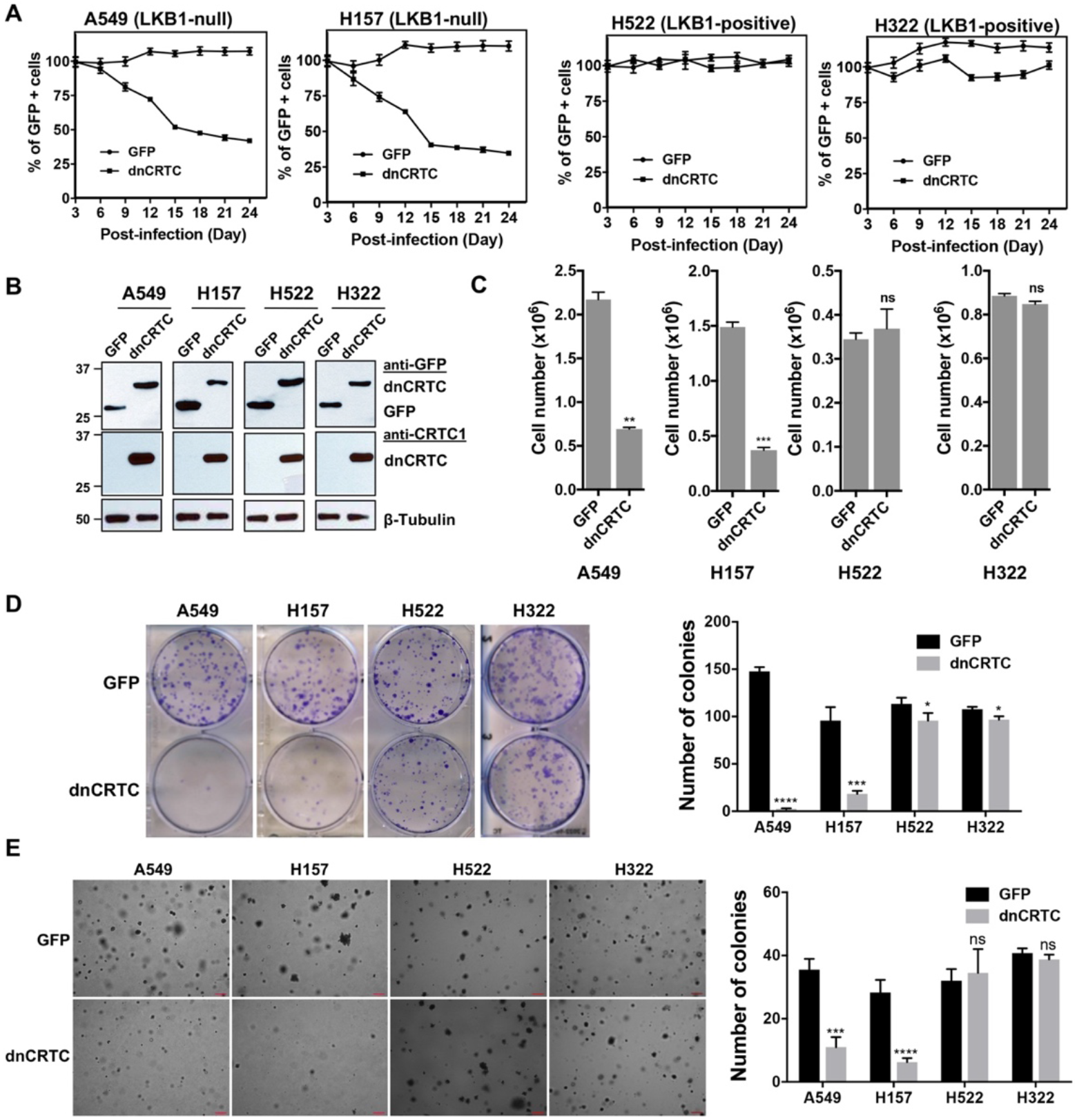
dnCRTC expression suppressed the growth of LKB1-null but not LKB1-positive lung cancer cells. **(A)** Two LKB1-null (A549 and H157) and two LKB1-positive (H322 and H522) NSCLC cells were transduced with dnCRTC or control GFP retroviruses. The MOI was optimized to obtain an infection rate of 40-60%, and then the percentage of GFP-positive cells was determined by FACS analysis every 3 days for a total of 24 days starting at day 3 post-infection. The percentage of GFP-positive cells at day 3 post-infection was considered as 100%, and the remaining data were normalized (n=3). **(B, C)** The GFP-positive cells for dnCRTC- and GFP-transduced cells were sorted and confirmed for dnCRTC and GFP expression by Western blotting (**b**). Sorted cells were also cultured at 2 ×10^5^ (for H322 and H522) or 3 × 10^5^ (for A549 and H157) cells/well in the 6-well plates for 96 hours and viable cells were counted using trypan blue exclusion test (**c**) (n=3). **(D)** Transduced cells were cultured at 400 cells/well in 6-well plates for 14 days and colonies were stained by crystal violet and photographed. The number of colonies in each well was counted using ImageJ. Assays were performed in triplicate (n=3). **(E)** Transduced cells were cultured in soft agar gels and colonies were stained by crystal violets, photographed and counted. The number of colonies from each image was counted using ImageJ. Assays were performed in triplicate. Scale bars, 200uM. Only colonies with a diameter higher than 50um were counted (n=3). Two-tailed student’s t-test was used to calculate the *p v*alues (*p<0.05, **p<0.01, ***p<0.001, ****p<0.0001, ns p>0.05).

### Inhibition of CRTC co-activators via dnCRTC expression blocked lung tumor growth and metastatic colonization *in vivo*

We further determined the effects of dnCRTC expression on the growth and metastatic colonization of lung cancer using subcutaneous and orthotopic NSCLC xenograft models. For subcutaneous xenograft models, luciferase-expressing LKB1-inactivated lung cancer cells (A549-luc and H157-luc) were transduced with retroviruses expressing dnCRTC or control GFP for 72 hours, and then dnCRTC or GFP-transduced cells (10^6^ cells per mouse) were subcutaneously implanted into immunodeficient NOD/SCID mice. The dnCRTC cohorts had reduced growth of xenograft tumors compared to the GFP control cohorts, as demonstrated by the reduced tumor growth rate, size and weight (**Figure 6A-D, F-I)**. Immunohistochemical analysis revealed a decreased number of Ki-67-positive proliferating cells in the dnCRTC-expressing xenograft tumors in comparison with the control GFP group (**Figure 6E, J**). Since the tumor cells used in these xenograft assays were unsorted and not 100% transduced, we performed Western blot analysis on the excised xenograft tumors and observed markedly reduced dnCRTC expression when compared to dnCRTC-transduced cells at the time of the injection. In contrast, GFP expression was similar between the excised GFP xenograft tumors and GFP-transduced cells at the time of injection (**Supplemental Figure 4**). These results indicate that the residual small xenograft tumors in the dnCRTC group were likely derived from cells with low or no dnCRTC expression, further supporting the tumor inhibitory effect of dnCRTC expression on the growth of LKB1-null lung cancers.

**Figure 6:**
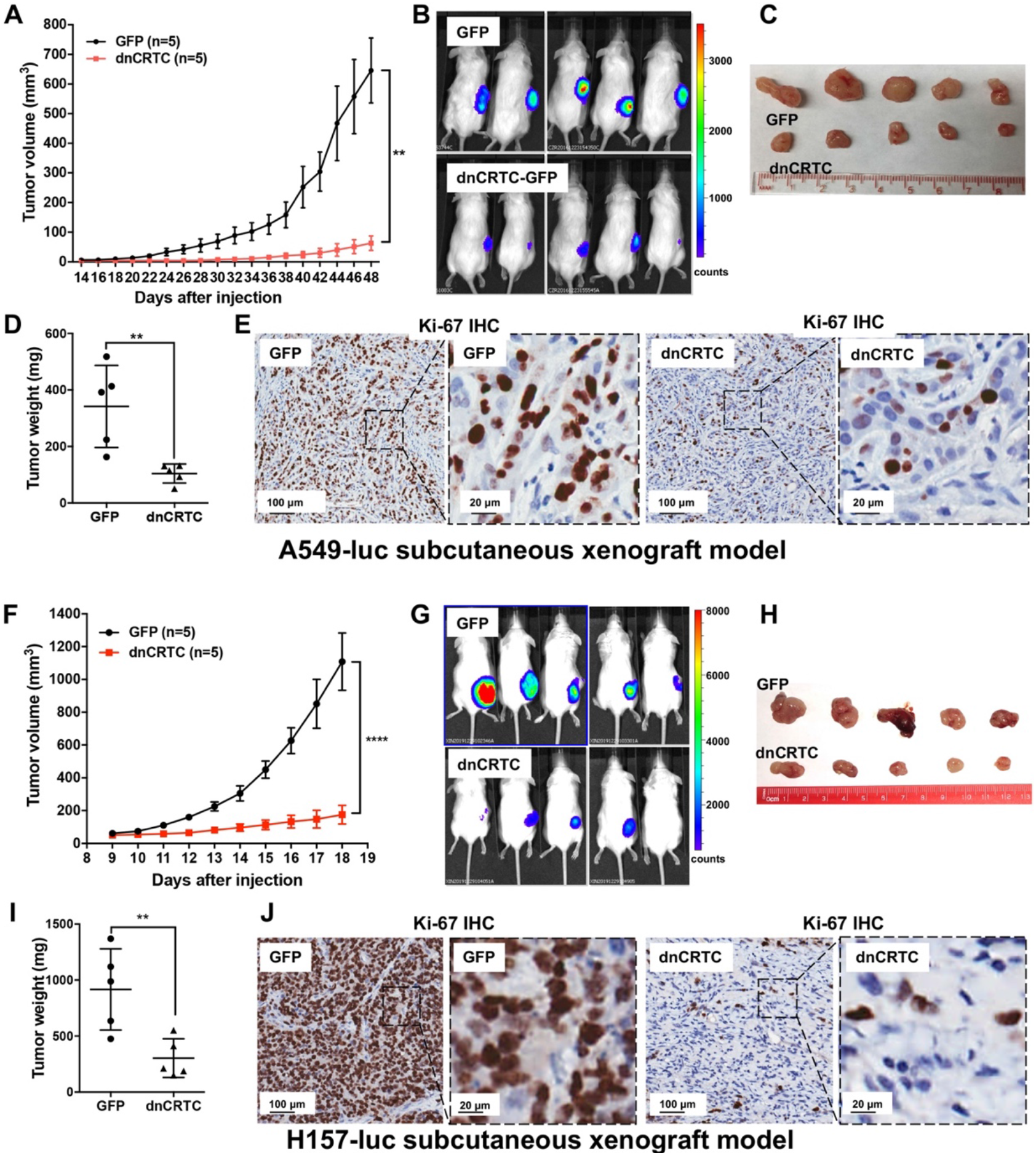
Expression of dnCRTC significantly inhibited the growth of LKB1-null NSCLC xenograft tumors. **(A-E)** A549-luc were transduced with GFP control or dnCRTC for 72 hours and the transduced cells (1×10^6^ per mouse) were injected subcutaneously to the right flanks of NOD/SCID mice. Tumor volumes of two cohorts (n=5 each) were measured every two days starting from day 14 until day 48 (A). The bioluminescent images of mice (B), excised tumors (C) and tumor weights (D) as well as Ki-67 immunohistochemical staining of xenograft tumor sections (E) were shown. **(F-J)** H157-luc were transduced with GFP control or dnCRTC for 72 hours and the transduced cells (1×10^6^ per mouse) were injected subcutaneously to the right flanks of NOD/SCID mice (n =5 each). Tumor volumes of two cohorts (n=5 each) were measured daily from day 9 to day 18 (F). The bioluminescent images of mice (G), excised tumors (H), tumor weights (I) and Ki-67 immunohistochemical staining (J) were shown. Scale bars: 100 μm (left panels), 20 μm (right panels). Two-tailed student’s t-test was used to calculate the *p v*alues (*p<0.05, **p<0.01, ****p<0.0001).

We also studied the effect of dnCRTC expression on the ability of lung cancer cells to undergo vascular extravasation and lung colonization using orthotopic NSCLC xenograft models. Here, dnCRTC-expressing or control GFP-expressing A549-luc cells or H157-luc cells (2×10^6^ per mouse) were intravenously injected into immunodeficient NOD/SCID mice and lung tumor burden was monitored. We observed that mice injected with dnCRTC-expressing A549-luc or H157-luc cells, compared to mice with their control GFP counterparts, had reduced tumor burden, a decreased number of surface tumor nodules and smaller tumor areas in the lung, as assessed by bioluminescent imaging **(Figure 7A, 7D)**, fluorescence imaging **(Figure 7B,7E)**, and H&E staining of lung sections (**Figure 7C, 7F**). Taken together, these data showed that expression of dnCRTC blocked lung cancer growth and colonization *in vivo*, indicating that the CRTC-CREB activation is essential for the growth and progression of LKB1-null lung cancer.

**Figure 7:**
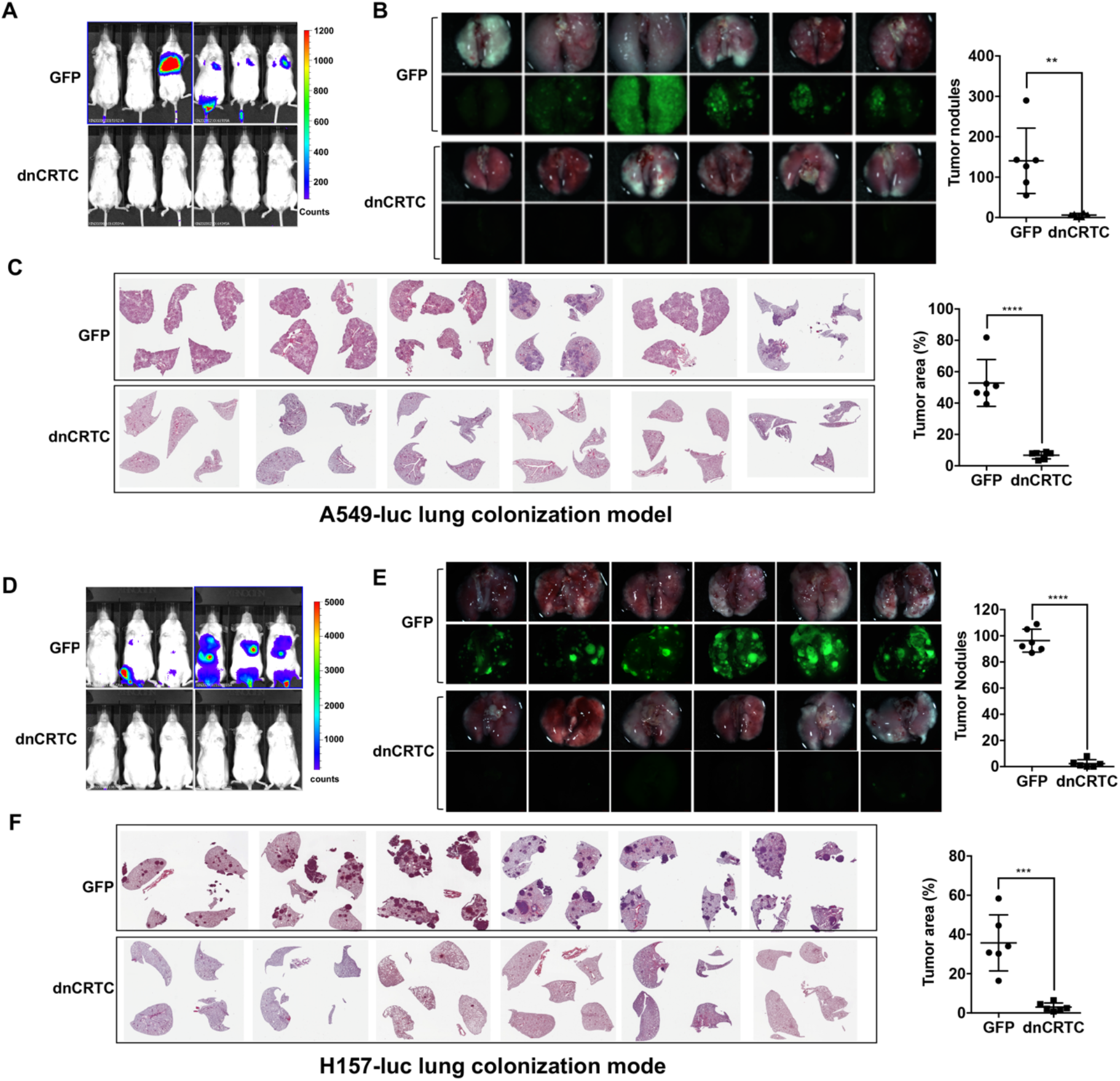
Expression of dnCRTC reduced lung colonization of LKB1-null lung cancer cells. **(A-C)** luciferase-expressing LKB1-null A549 cells (A549-luc) were transduced with retroviruses expressing GFP control or dnCRTC for 72 hours, and transduced cells (2×10^6^ cells per mouse) were intravenously injected to NOD/SCID mice (n=6 each). Eight weeks after injection, lung colonization was assessed by bioluminescent imaging (A). Lungs were dissected and bright field and GFP fluorescence images were shown (B). The number of surface tumor nodules with visible GFP signal per lung of each mouse was quantified and presented (right panel). Representative H&E staining of lung sections were shown (C). Tumor area was calculated from multiple H&E-stained lung sections from each mouse and presented as a percentage of tumor area to total lung area (right panel). **(D-F)** Luciferase-expressing LKB1-null H157 lung cancer cells (H157-luc) were transduced with retrovirus expressing GFP control or dnCRTC for 72 hours, and transduced cells (2×10^6^ cells per mouse) were intravenously injected to NOD/SCID mice (n=6 each). Four weeks after injection, lung colonization was assessed by bioluminescent imaging (D). Lungs were dissected and bright field and GFP fluorescence images were shown (e). The number of tumor nodules with visible GFP signal per lung of each mouse was quantified (right panel). Representative H&E staining images of lung sections were shown (F). Tumor area was calculated from multiple H&E-stained lung sections from each mouse and presented as a percentage of tumor area to total lung area (right panel). The p values were calculated by two-tailed student’s t-test (**p<0.01, ***p<0.001, ****p<0.0001).

## Discussion

Lung cancer carrying somatic *LKB1* inactivation is a common aggressive molecular subtype with very limited treatment options. Since replacing loss-of-function tumor suppressor mutations is challenging, drug therapeutic efforts have been directed towards identifying and understanding the effector pathways that mediate *LKB1* tumor suppression in order to uncover new therapeutic strategies. An important function of LKB1 is its ability to activate SIKs which then phosphorylate and negatively regulate the family of three CREB-regulated transcriptional co-activators (CRTC). We and others have shown that the loss of LKB1 directly leads to CRTC activation and extensive, elevated CRTC1-CREB-mediated transcription in human lung cancer cells and primary tumors (33–35, 41, 42, 46). More recently, studies of genetically engineered mouse models of oncogenic *KRAS*-induced lung cancer revealed that SIKs, but not other AMPK family members, mediate the major tumor suppression function of LKB1 (40, 41). These molecular and genetic data support the model that aberrant CRTC-CREB transcriptional activation mediates the major LKB1-null malignancy. However, direct evidence for the importance of CRTC activation in promoting tumorigenesis was lacking. Also, whether there is a specific role for individual CRTC 1-3 family members was unknown. In this study, we showed overlapping expression and the potential for functional redundancy of three CRTC co-activators in lung cancers. Therefore, we designed and validated a pan-CRTC dominant negative inhibitor as a useful tool for blocking all three CRTC co-activator function. Our new mechanistic and functional data demonstrated an essential, general role for CRTC activation in maintaining the malignant phenotypes of LKB1-inactivated lung cancer and identified the CRTC-CREB interaction as a valuable molecular target for development of new therapies for lung cancer with LKB1 deficiency.

The findings in this study further emphasize the importance of CRTC activation in tumorigenesis. We initially identified CRTC1 as a fusion partner with the Notch transcriptional co-activator MAML2, due to a t(11;19) chromosomal translocation in mucoepidermoid carcinoma (MEC), the most common salivary gland malignancies and lung tumors (39, 47, 48). This fusion event leads to a chimeric CRTC1-MAML2 protein which is composed of the CREB-binding domain (CBD) of CRTC1 (42aa) fusing to the transcriptional activation domain (TAD) of MAML2 (983aa) (39, 48). The CRTC1-MAML2 fusion binds to CREB via the CRTC1 CBD and potently activates CREB-dependent transcription through its MAML2 TAD (48–50), which contribute to the fusion’s major oncogenic activity. These data demonstrate a critical role of CRTC activation in MEC tumorigenesis. In our previous studies, we also showed that LKB1-deficiency led to CRTC activation of many CREB-dependent genes, including NR4A2, PTGS2 (aka COX-2), LYPD3, INSL4 and LINC00473, which play important roles in cancer cell growth, survival or invasive properties (34, 35, 51, 52). Recently, other groups reported that SIKs were the major AMPK family members that mediate LKB1 tumor suppression and SIK knockouts enhanced CRTC target gene expression (41, 46). Furthermore, CRTC2 downregulation inhibited the growth of LKB1-deficient NSCLC (42). All these data support a new model that LKB1-SIK genetic alterations represent a distinct mechanism for the constitutive CRTC-CREB activation that is critical for the tumorigenesis and progression of NSCLC. In this study, we performed expression and functional assays to determine the relative contributions of three CRTC co-activators in lung cancer cells. Our data showed that all three CRTC co-activators (CRTC1-3) are expressed at various levels in lung cancers and that CRISPR/Cas9-mediated knockouts of individual CRTCs only partially reduced the LKB1 target gene expression and had very moderate impact on lung cancer cell proliferation, colony formation and anchorage-independent growth. It should be noted that a recent study reporting that CRTC2 shRNA knockdown or knockout in polyclonal cells impaired soft agar formation but did not affect cell proliferation (42). In our study, two CRTC2 KO single clones only showed minimal inhibition of cell proliferation and soft agar colony formation; this discrepancy could be explained by the upregulation of CRTC1 in the CRTC2 KO clones. Therefore, these data indicate the presence for functional redundancy of the three CRTC co-activator family members in driving aberrant CREB transcriptional program and lung cancer malignant phenotypes, thus suggesting that general inhibition of all CRTCs is required for blocking the aberrant CRTC-induced transcriptional program and lung tumorigenesis.

We subsequently developed a dominant negative mutant dnCRTC to block all 3 CRTC function. This dnCRTC binds to CREB but is defective in transcriptional activation, consequently forming an inactive transcriptional complex with CREB and interfering with the ability of all three CRTCs to co-activate CREB-mediated transcription. Expression of this pan-CRTC inhibitor efficiently and extensively inhibited the aberrantly activated CREB-mediated transcriptional program induced by LKB1 deficiency. Integrated analysis of the dnCRTC-regulated DEGs from our gene expression profiling with the published ChIP-seq data (42) revealed the direct target genes downstream of CRTC activation, which include known and potential novel mediators of aberrant CRTC activation in LKB1-inactivated cancer. Future studies of these mediators of CRTC activation and their potential cross-talk with other signaling pathways will enhance our molecular understanding of the loss-of-LKB1 tumor suppression in lung cancer. Since dnCRTC acts as a pan-CRTC inhibitor, it has the potential to serve as an invaluable research tool for dissecting the role of deregulated CRTC activation in various disease settings, such as cancers with aberrant CRTC activation (e.g. LKB1 deficiency, the CRTC1-MAML2 fusion), diabetes with CRTC activation that contributes to high blood glucose levels as well as neurological conditions such as depression and memory.

In this study, we utilized this pan-CRTC inhibitor to probe the functional impact of blocking the CRTC-CREB activation on the growth of multiple NSCLC cell lines and xenograft models. We showed that dnCRTC expression caused significant growth inhibition in LKB1-null, but not LKB1-wt cancer cells and normal lung epithelial cells. The growth and lung colonization of LKB1-null lung cancer cells were specifically susceptible to inhibition of CRTC coactivators. These results demonstrate an essential role of aberrant CRTC activation in supporting the malignant phenotypes of LKB1-inactivated lung cancers. Future research examining the effect of CRTC inhibition in lung cancer genetic mouse models is needed to elucidate the role of CRTC co-activators in lung cancer development and progression.

As the malignant phenotypes of LKB1-inactivated lung cancer are specifically dependent on aberrant CRTC co-activation of the oncogenic CREB transcriptional program, targeting the CRTC-CREB interaction, hence, the active CRTC-CREB transcription complex, may selectively inhibit LKB1-deficient tumors with minimal effects on normal cells as demonstrated by our data. The strategy of blocking the assembly of active CRTC-CREB transcriptional complex and subsequently inhibiting extensive CRTC target genes has the advantage of simultaneously inhibiting multiple deleterious cell signals which predicts a greater challenge for resistant cancer cell clones to emerge. We propose that inhibition of the CREB-CREB interaction should reverse the oncogenic activity of CRTC activation. For instance, peptides and peptide-like molecules designed to recapitulate a critical interaction motif will have the potential in selectively targeting the CRTC/CREB interaction interface and consequently inhibit lung cancer growth. The crystal structural analysis has provided important molecular information of the core CRTC/CREB transcriptional complex (53, 54), revealing that CRTC CBD interacts with CREB basic leucine zipper (bZIP) domain forming a 2:2 complex on CRE-containing DNA. CRTC interacts with both CREB and DNA through highly conserved residues are crucial for the complex assembly and CREB stabilization on DNA. With insights from the crystal structural studies and further understanding of the assembly and composition of the CRTC/CREB transcriptional complex, new approaches can be developed to inhibit the oncogenic CRTC/CREB transcriptional program and block the progression of lung cancers.

Collectively, our study provides direct proof for a critical role of the CRTC-CREB activation in maintaining the malignant phenotypes of LKB1-inactive lung cancers and reveals direct inhibition of the CRTC-CREB transcriptional complex via targeting the CRTC-CREB interface as a novel, promising therapeutic approach.

## Materials and methods

### Cell culture

Human NSCLC cancer cell lines (A549, H157, H322, H522, H2126, H1819, H2087, H2009, and H3123) were cultured in DMEM (Corning #10-013-CV) supplemented with 10% (vol/vol) heat-inactivated fetal bovine serum (Gibco #10437028), and penicillin (100 U/mL)/streptomycin (100 μg/mL) (HyClone #SV30010). Human NSCLC cancer cell lines (H23, H460, H2122 and H358) and mouse lung squamous carcinoma mLSCC^LP^ cell line (45) were cultured in RPMI-1640 (HyClone # SH3002701) with 10% inactivated fetal bovine serum and penicillin/streptomycin. Immortalized human bronchial epithelial BEAS-2B cells were purchased from ATCC and cultured in BEGM bronchial epithelial cell growth medium (Lonza #CC-4175). All the cells were grown at 37°C with 5% CO_2_.

### Plasmids

The sgRNA sequences targeting CRTC1, CRTC2, and CRTC3 were designed using the CRISPR design tool (https://zlab.bio/guide-design-resources) and cloned into the lentiCRISPR v2 vector that co-expresses Cas9 (#52961) (55). The control plasmid sgCtr-LentiCRISPRv2 expressing a non-target sgRNA was purchased from Addgene (#107402) (56). The sgRNA target sequences and non-targeting control were listed in Supplementary Table 2. The pMSCV-dnCRTC retroviral construct was generated by cloning a DNA fragment encoding the CRTC1 CBD domain (1-55 aa) followed by a nuclear localization signal (PKKKRKV) into the backbone of the pMSCV-GFP vector (57) by replacing the internal ribosome entry sequence (IRES). The cAMP response element (CRE) luciferase reporter (pCRE-luc), Renilla luciferase plasmid (pEF-RL), and pFLAG-CMV2 vectors expressing individual CRTC were previously described (48). The pcDNA FLAG-tagged CRTC2 (#22975) and FLAG-tagged CRTC3 (22976) expression constructs were obtained from Addgene (38).

### CRISPR-Cas9-mediated gene knockout

LentiCRISPR constructs containing sgRNAs for CRTC1, CRTC2, or CRTC3 or control sgRNA (Supplementary Table 2) were transfected into 293FT cells together with packaging plasmids pMD2.G and pSPAX2 using Effectene transfection reagent (Qiagen #301425). The viral supernatants were collected at 48, 72 and 96 hours after transfection. A549 cells were then infected by culture-medium-diluted viral supernatants in the presence of 6 μg/ml polybrene (Sigma #H9268) in 3 consecutive days and selected with puromycin (1.5 μg/ml) for 48 hours. Single-cell cloning was set up through serial dilutions in 96-well plates, followed by expansion of cell culture. The knockout clones were validated for the loss of protein expression by Western blotting and for altered genomic sequences by DNA sequencing.

### Retroviral transduction

293FT cells were transfected with pMSCV-dnCRTC or pMSCV-GFP constructs together with packaging plasmid pMD.MLV and pseudotyped envelope plasmid pMD2-VSV-G using Effectene transfection reagent (Qiagen #301425) as previously described. Viral supernatants were collected at 48 and 72 hours post-transfection. Targeted cells (A549, H157, H322, H522, BEAS-2B, mLSCCLP) were infected with viral supernatants mixed with fresh complete medium plus 6 ug/ml polybrene (Sigma #H9268) for 6 hours. Infection was performed twice in two consecutive days.

### Quantitative RT-PCR

Total RNA was isolated using RNeasy Mini Kit (Qiagen #74106) and then reverse-transcribed into complementary DNA using a High Capacity cDNA Reverse Transcription Kit (Applied Biosystems #4368814). PCR was subsequently performed using StepOne Real-Time PCR System with iTaq Universal SYBR Green Supermix (Bio-Rad #1725120). The relative gene expression was calculated using the comparative ∆∆Ct method. Glyceraldehyde-3-phosphate dehydrogenase (GAPDH) was used as an internal control for normalizing gene expression among different samples. The primer sequences were listed in Supplementary Table 2.

### Western blotting analysis

Cells were lysed in lysis buffer [10 mM Tris/Cl pH 7.5, 150 mM NaCl, 0.5 mM EDTA, 0.5% NP-40, 2 mM Na3VO4, 1mM PMSF, 2 mg/ml protease inhibitor cocktail (cOmplete, Roche)] on ice for 30 minutes. Protein lysates were collected after removing insoluble fractions by centrifugation at 13000 rpm for 15 minutes at 4℃. Protein lysates (~50 μg/lane) were separated on 7.5% SDS-PAGE gels and electrophoretically transferred onto nitrocellulose membranes. The membranes were blocked in 5% w/v fat-free milk in TBST buffer (10 mM Tris-HCl, pH 8.0, 150 mM NaCl, 0.05% Tween 20) at room temperature for one hour and then incubated with primary antibodies diluted in TBST at 4°C overnight. After extensive washing with TBST, the membranes were incubated with horseradish peroxidase (HRP)-coupled secondary antibodies at room temperature for one hour, washed again and proteins were visualized by SuperSignal™ West Dura Extended Duration Substrate (Qiagen #34076)

The following antibodies were used for Western blotting: anti-CRTC1 (Cat #600-401-936, Rabbit) from Rockland Immunochemicals Inc; anti-CRTC2 (Cat #A300-637A, Rabbit) and anti-PDE4D (Cat #A302-744A, Rabbit) from Bethyl Laboratories; anti-CRTC3 (Cat #2720, Rabbit), anti-LKB1 (Cat #3050, Rabbit) from Cell Signaling Technology; anti-β-TUBULIN (Cat #1878, Rabbit) from Epitomics; and anti-β-ACTIN (Cat #A5316, Mouse) from Sigma-Aldrich.

### Chromatin immunoprecipitation

ChIP assay was performed based on a ChromoTek GFP-Trap immunoprecipitation protocol (ChromoTek #gtma-10). The dnCRTC- or GFP-expressing A549 cells were fixed with 1% formaldehyde for 10 minutes at room temperature and lysed using lysis buffer (10 mM Tris/Cl pH 7.5; 150 mM NaCl; 0.5 mM EDTA; 0.5% NP-40; 1mM PMSF). The chromatin was sonicated to around 100-500bp. The DNA-protein complex was then immunoprecipitated with GFP-trap or control IgG agarose beads, and analyzed for CREB and dnCRTC by Western blot. The ChIP DNA was also purified and used for real-time PCR assays using the primers that amplify the regions spanning the CRE sites of LINC00473 and NR4A2 promoters. The primer sequences were list in Supplementary Table 2.

### Luciferase reporter assays

HEK293T cells were seeded in 24-well plates at 1×10^5^ cells/well overnight and transfected with pCRE-luc firefly luciferase vector, internal control Renilla luciferase plasmid (pEF-RL), pFLAG-CMV2 vectors expressing CRTC1, or CRTC2, or CRTC3, and pMSCV-GFP or pMSCV-dnCRTC using Effectene transfection reagent (Qiagen #301425). The luciferase assays were carried out at 48 hours after transfection using a dual-luciferase assay kit (Promega, #E1910) as described previously (48).

### Transcriptomic analysis

Two biological replicates of RNA samples were isolated from A549 cells transduced with pMSCV-dnCRTC or pMSCV-GFP retroviruses at 72 hours post-infection and were then subjected to microarray experiment using GeneChip® Human Transcriptome Array 2.0 (Affymetrix) at the Genomics Core at Sanford Burnham Research Institute. The same samples were also subjected to RNAseq analysis by Novogene. In brief, RNA-seq libraries (non-strand-specific, paired end) were prepared with the NEBNext^®^ Ultra™ RNA Library Prep Kit (Illumina) and were sequenced according to the paired-end 150bp protocol using NovaSeq 6000. The data were analyzed as previously described (33, 34, 50). Genes with an absolute fold change ≥2 and an FDR p-value <0.5 were considered as significantly differentially expressed. The data were deposited in the NCBI GEO database (GSE157722).

### Cell growth competition assay

The competitive cell growth assay was performed as previously described (48). In brief, lung cancer cells (A549, H157, H322, H522) were infected with pMSCV-based retroviruses expressing dnCRTC or GFP at infection rates between 40%-60%. Cells were seeded and harvested every 3 days. The percentage of GFP-positive cells was determined by flow cytometry every 3 days for a total of 24 days. The percentage of GFP-positive cells at day 3 after the infection was considered as 100%, and the remaining data were normalized.

### Cell proliferation and apoptosis assays

Cells were seeded in the 6-well plates at 0.3×10^6^ cells/well and cultured for 96 hours for cell proliferation and apoptosis assays. Cell proliferation was determined by direct counting of viable cells stained with 0.2% trypan blue solution. For apoptosis assay, cells were stained using a FITC Annexin V Apoptosis Detection Kit (BD Bioscience Cat #556547) and analyzed by Accuri C6 Flow Cytometer (BD Biosciences). Cells with Annexin V positive and PI positive or negative were considered apoptotic cells.

### Colony formation and soft agar assays

For colony formation assay, cells were grown in 6-well plates at 400 cells per well for 14 days. Each well was then fixed by fixation buffer (12.5% acetic acid/87.5% methanol) for 30 minutes followed by staining with 0.1% crystal violet solution (0.1% crystal violet, 10% ethanol dissolved in ddH_2_O) for another 30 minutes at room temperature. Plates were then washed with tap water, air-dried and scanned. The number of colonies from each well was counted using ImageJ. Assays were performed in triplicate.

Soft agar assays were performed in 6-well plates with 20,000 cells per well. Cells were suspended as single cell suspension in culture medium containing 0.35% noble agar (BD Biosciences, #214230) and then layered on the top of 0.5% agar in culture medium). The plates were incubated for 14 days and were stained by crystal violet solution (0.5% crystal violet-10% ethanol) at room temperature and colonies photographed under a microscope. Four images in different fields for each well were obtained. The number of colonies from each image was counted using ImageJ. Only colonies with a diameter higher than 50um were counted. Assays were performed in triplicate.

### Mouse xenograft assay

For subcutaneous xenograft assay, A549 and H157 cells stably expressing firefly luciferase (A549-luc/H157-luc) were infected with pMSCV-dnCRTC or pMSCV-GFP retroviruses. A total of 1×10^6^ cells were diluted in 100ul 50% Matrigel (BD Biosciences) and injected subcutaneously to dorsal flanks of 8-12 week-old NOD.SCID mice (Jackson Laboratory; stock# 001303). Tumors were measured using a vernier caliper every 1-2 days and tumor volumes were calculated using the formula: tumor volume = (length × width^2^) ×0.5. At the endpoint, mice were given 150 mg/g of D-luciferin in PBS by intraperitoneal injection for 10 minutes and bioluminescence was then imaged with a Xenogen In-vivo Imaging System (Caliper Life Sciences). Mice were then euthanized and tumors were dissected, photographed, weighed, fixed in 4% paraformaldehyde at 4°C for 48 hours and embedded into paraffin blocks.

For orthotopic xenograft assay, luciferase-expressing A549 and H157 cells (A549-Luc/ H157-Luc) were transduced with retroviruses containing pMSCV-dnCRTC or pMSCV-GFP. A total of 2×10^6^ cells were diluted in 100μl PBS and intravenously injected into NOD/SCID mice from the tail vein. At the endpoint, bioluminescence was imaged as described above. Mice were euthanized and perfused with PBS. Lungs were then dissected out and photographed under a fluorescence stereomicroscope (Leica MZ16 F). The number of tumor nodules on the lungs in each mouse were counted under the microscope. Lungs were fixed in 4% paraformaldehyde at 4°C for 48 hours and paraffin embedded. Paraffin tissue sections with 4 uM thickness were prepared and H&E staining and Ki-67 IHC staining were performed at the Molecular Pathology Core, University of Florida (Gainesville, FL) as previously described (33). Total tumor burden (tumor area/total area ×100%) was quantified from H&E sections using ImageJ.

### Statistics

Data were analyzed using GraphPad Prism 7 (GraphPad Software, Inc., USA). The statistical significance was determined by two-tailed Student’s *t*-test for two groups or by one-way ANOVA test for multiple groups (>2). Results were presented as the mean ± SD, and *p*-value < 0.05 was considered statistically significant.

### Study approval

Animal studies were performed following a protocol approved by the IACUC (Institutional Animal Care & Use Committee) of the University of Florida.

## Authors’ contributions

X.Z. performed experiments, analyzed data, and wrote the manuscript;

J.W.L: performed data analysis, and edited the manuscript;

Z.C., W.N., X.L., R.Y.: performed experiments;

H.S., J.L., F.J.D., J.L., F.J.K: provided reagents and suggestions, and assisted with data interpretation and manuscript writing;

L.W.: designed and directed the study, analyzed and interpreted data, and wrote the manuscript. All authors read and approved the final manuscript.

## Acknowledgments

We thank the ICBR Cytometry Core and Molecular Pathology Core at the University of Florida for the technical support. This work was supported by the National Institutes of Health (R01CA234351 and R01DE023641 to LW.; Z1AES103311-01 to FJD), the University of Florida Gatorade Trust (to FJK), and UF Health Cancer Center.

