## Supplemental Information for "Dependency of LKB1-inactivated lung cancer on aberrant CRTC-CREB activation"

**Supplemental Figures: S1-S4**

Figure S1: Page 2

Figure S2: Pages 3-4

Figure S3: Page 5

Figure S4: Page 6

**Supplemental Tables: S1-2**

Table S1: Pages 7-12

Table S2: Pages 13

### Supplemental Figure 1

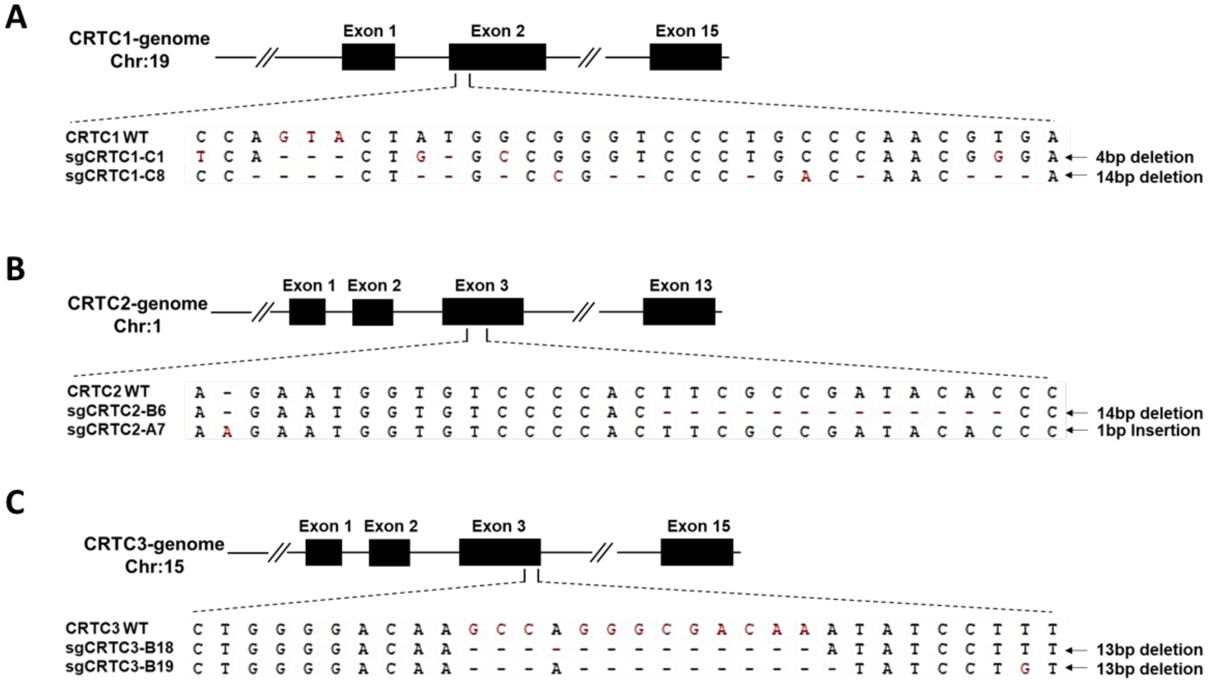


**Figure S1: CRISPR/Cas9-edited alleles in two independent single knockout clones for each CRTC gene were validated by genomic DNA sequencing.** The altered sequences of *CRTC1* (A), *CRTC2* (B) and *CRTC3* (C) genes in two independent single knockout clones were shown.

### Supplemental Figure 2

### A

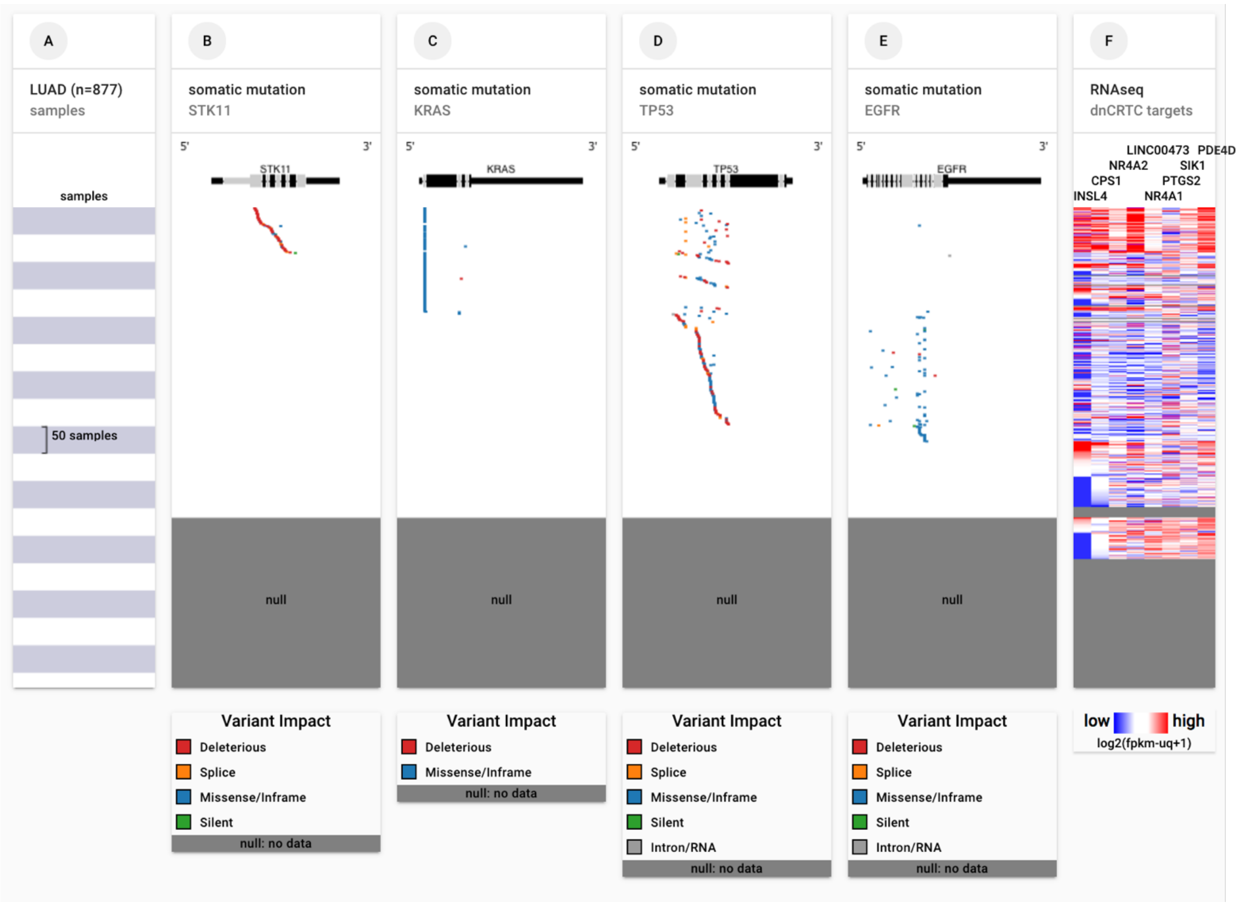

**B**

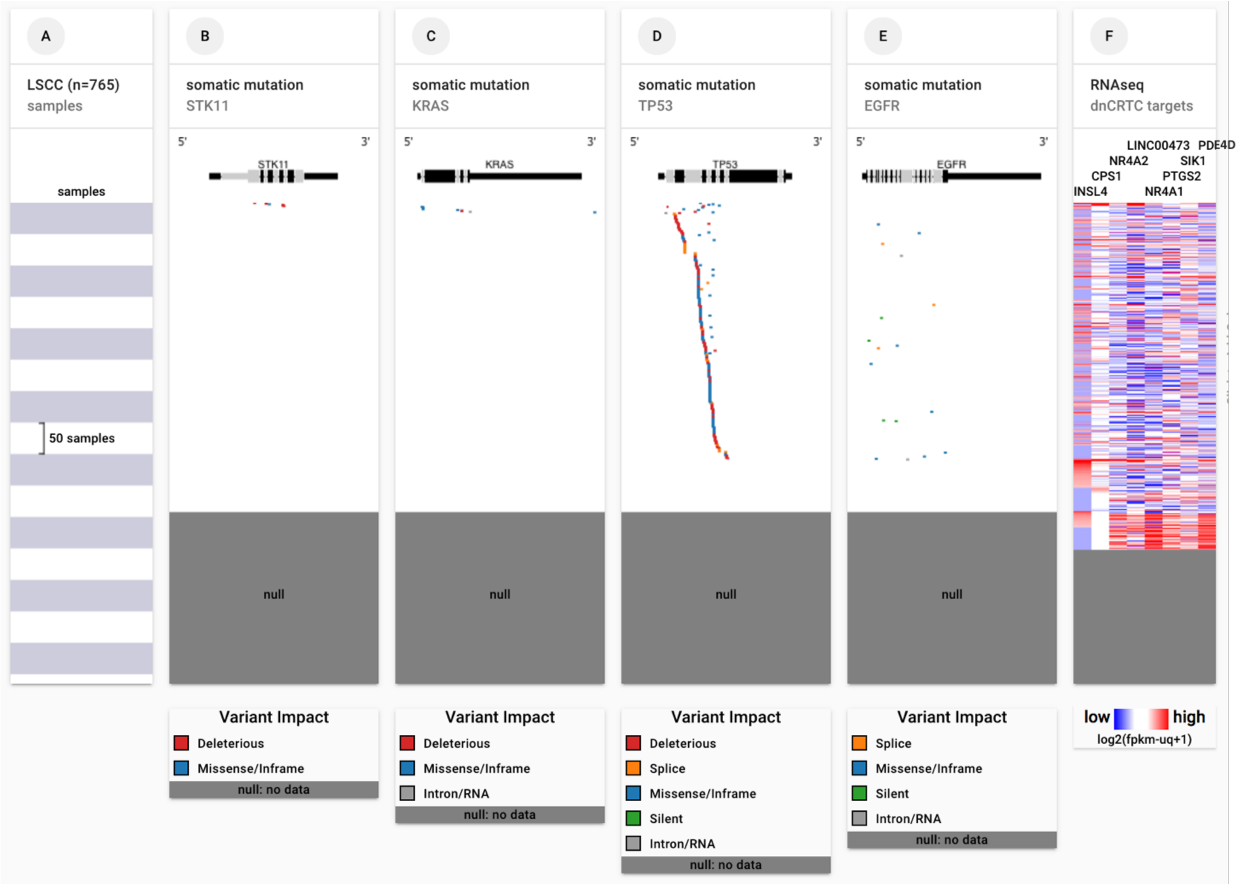


**Figure S2: The high expression levels of multiple dnCRTC target genes were associated with LKB1 mutations in human NSCLC tumors.** The overall high expression of several top dnCRTC target genes, such as *INSL4*, *CPS1*, *NR4A2*, *LINC00473*, *NR4A1*, *PTGS2*, *SIK1*, *and PDE4D,* was associated with somatic mutations (SNPs and small INDELs) in *LKB1*, but not *KRAS*, *TP53* or *EGFR*, in both human lung adenocarcinoma (LUAD) **(A)** and lung squamous cell carcinoma (LSCC) **(B)** cohorts. The data were generated by UCSC Xena using the genomic and RNAseq data of the GDC TCGA LUAD and LSCC cohorts.

### Supplemental Figure 3

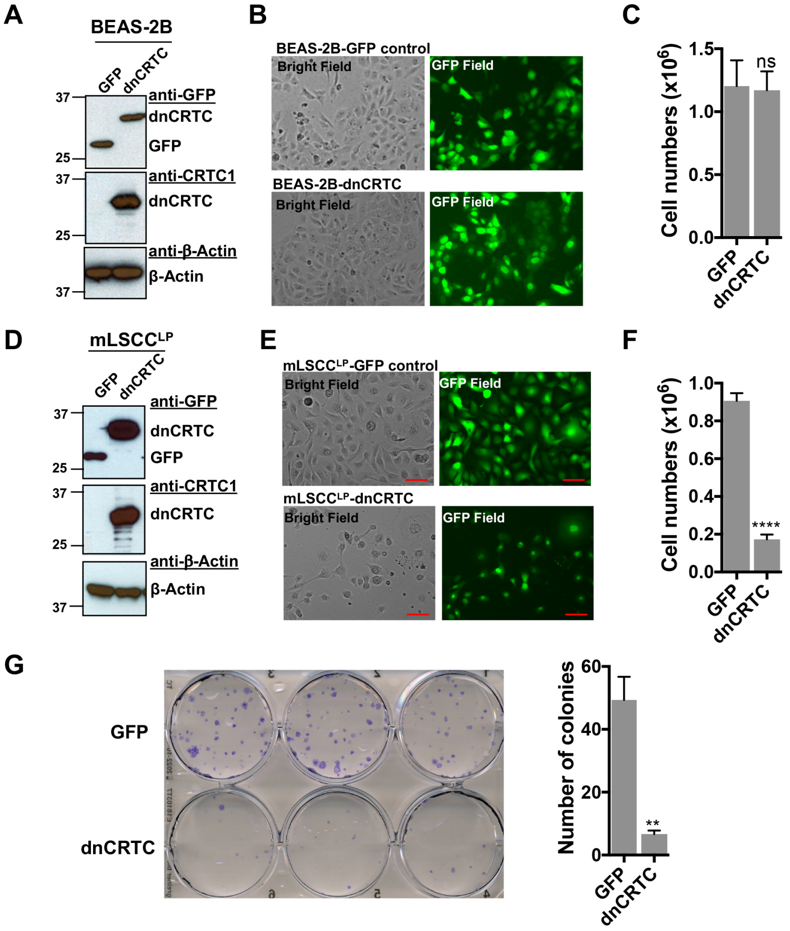


**Figure S3: Effects of dnCRTC expression on the growth of human immortalized lung bronchial epithelial BEAS-2B cells and mouse LKB1-null NSCLC mLSCC$^{LP}$ cells. (A)** BEAS-2B cells were transduced with dnCRTC or GFP retroviruses and the dnCRTC and GFP expression were confirmed by Western blotting. **(B)** The transduced dnCRTC- or GFP BEAS-2B cells were photographed under a microscope, and cell images under both the bright field and GFP field were presented. **(C)** The BEAS-2B cells transduced with dnCRTC or GFP retroviruses were cultured at $3x10^5$ cell per well in 6-well plates for 96 hours. The number of viable cells were determined by trypan blue exclusion assay (n=3). **(D)** Mouse NSCLC mLSCC$^{LP}$ cells with LKB1 and PTEN deficiency were transduced with dnCRTC or GFP retrovirus. The dnCRTC and GFP expression were confirmed by Western blotting. **(E)** The transduced dnCRTC- or GFP mLSCC$^{LP}$ cells were photographed and cell images under both the bright field and GFP field were presented. **(F)** mLSCC$^{LP}$ cells transduced with dnCRTC or GFP-control retroviruses were cultured at $2X10^5$ cell per well in 6-well plates for 96 hours. The number of viable cells were determined by trypan blue exclusion assay (n=3). **(G)** The transduced mLSCC$^{LP}$ cells were cultured at 400 cells per well in 6-well plates for 14 days and the colonies were stained by crystal violet and photographed. The number of colonies in each well was counted using ImageJ. Assays were performed in triplicate. The p values were calculated by two tailed student's t-test (**p<0.01, ****p<0.0001, ns p>0.05)

### Supplemental Figure 4

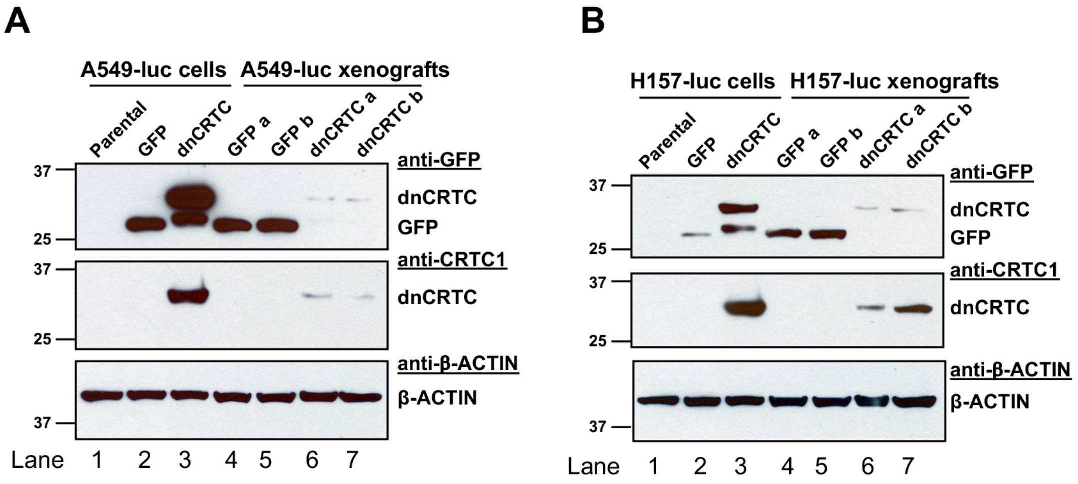


**Figure S4: Expression of GFP and dnCRTC in the transduced human LKB1-null lung cancer cells at the time of tumor cell injection and in the resulting xenograft tumors**. **(A)** Western blot analysis of GFP and dnCRTC proteins in luciferase-expressing A549 (A549-luc) cells and xenograft tumors. Lane 1: A549-luc (parental); Lane 2: A549-luc transduced with GFP control retroviruses for 72 hours; Lane 3: A549-luc transduced with dnCRTC retroviruses for 72 hours; Lane 4-5: 2 xenograft tumors derived from 2 mice implanted with GFP-expressing A549-luc cells; Lane 6-7: 2 xenograft tumors derived from 2 mice implanted with dnCRTC-expressing A549-luc cells. **(B)** Similar Western blotting analysis was performed as (A) except that H157-luc cells were used.

**Table S1: Differentially expressed genes in dnCRTC-expressing A549 lung cancer cells in comparison with GFP-expressing control cells were shown.** The cutoff criteria included an asolute fold change >= 2.0 and FDR <=0.05. Genes with the predicted CREB binding sites based on the CREB Target Gene Database (Ref 44), or CREB binding or CRTC2 binding from a ChIP-seq study (Ref 42) were shown.

| Probe Set ID | Gene Accession | Gene Symbol | Gene Description | Fold Change (dnCRTC/GFP) | FDR (dnCRTC vs GFP) | Predicted CREB site (-3kb to 300bp) | CREB binding (-3kb to 300bp) | CREB binding (-500b to 100bp) | CRTC2 binding (-3kb to 300bp) | CRTC2 binding (-500b to 100bp) |
|---|---|---|---|---|---|---|---|---|---|---|
| TC09000036.hg.1 | NM_002195 | INSL4 | insulin-like 4 (placenta) | -218.82 | 9.16E-07 | CRE | | | Yes | Yes |
| TC02001246.hg.1 | NM_001122633 | CPS1 | carbamoyl-phosphate synthase 1, mitochondrial | -33.50 | 6.05E-09 | CRE | | | | |
| TC02002445.hg.1 | NM_006186 | NR4A2 | nuclear receptor subfamily 4, group A, member 2 | -26.36 | 8.36E-09 | CRE | Yes | Yes | Yes | Yes |
| TC07001618.hg.1 | NM_002612 | PDK4 | pyruvate dehydrogenase kinase, isozyme 4 | -25.87 | 7.65E-09 | CRE | | | Yes | Yes |
| TC06002302.hg.1 | NR_026860 | LINC00473 | long intergenic non-protein coding RNA 473 | -23.63 | 1.01E-06 | CRE | Yes | Yes | Yes | Yes |
| TC12000414.hg.1 | NM_001202233 | NR4A1 | nuclear receptor subfamily 4, group A, member 1 | -18.78 | 6.36E-08 | CRE | Yes | Yes | Yes | Yes |
| TC02002835.hg.1 | NM_024795 | TM4SF20 | transmembrane 4 L six family member 20 | -17.32 | 1.14E-07 | | | | | |
| TC09000508.hg.1 | NM_006981 | NR4A3 | nuclear receptor subfamily 4, group A, member 3 | -16.47 | 3.94E-07 | CRE | Yes | Yes | Yes | Yes |
| TC02001247.hg.1 | NR_002763 | CPS1-IT1 | CPS1 intronic transcript 1 (non-protein coding) | -10.27 | 1.02E-05 | | | | | |
| TC01003638.hg.1 | NM_000963 | PTGS2 | prostaglandin-endoperoxide synthase 2 (prostaglandin G/H synthase an | -8.20 | 2.36E-05 | CRE | Yes | | Yes | |
| TC21000506.hg.1 | NM_173354 | SIK1 | salt-inducible kinase 1 | -7.33 | 1.62E-06 | CRE | | | | |
| TC07000562.hg.1 | NM_001040152 | PEG10 | paternally expressed 10 | -6.82 | 1.10E-06 | CRE | | | | |
| TC10001569.hg.1 | NM_021732 | AVPI1 | arginine vasopressin-induced 1 | -6.52 | 1.30E-05 | CRE | Yes | Yes | Yes | Yes |
| TC08001692.hg.1 | OTTHUMT000003 | RP11-10J21.4 | novel transcript | -6.02 | 2.34E-05 | | | | | |
| TC01000757.hg.1 | NM_001190463 | CTH | cystathionine gamma-lyase | -5.84 | 9.16E-07 | CRE | Yes | Yes | Yes | Yes |
| TC01001607.hg.1 | NM_024420 | PLA2G4A | phospholipase A2, group IVA (cytosolic, calcium-dependent) | -5.80 | 1.01E-06 | CRE | | | | |
| TC06002300.hg.1 | ENST0000053522 | PDE10A | phosphodiesterase 10A | -5.59 | 0.0006168 | CRE | | | Yes | |
| TC04000777.hg.1 | NM_001184741 | FGB | fibrinogen beta chain | -5.45 | 7.39E-06 | CRE | | | | |
| TC04001382.hg.1 | NM_198281 | GPRIN3 | GPRIN family member 3 | -5.43 | 1.12E-05 | CRE | | | | |
| TC10000968.hg.1 | ENST0000045946 | RNU7-163P | RNA, U7 small nuclear 163 pseudogene | -5.29 | 0.0016756 | | | | | |
| TC12002012.hg.1 | NM_001168325 | TESC | tescalcin | -5.25 | 6.94E-07 | CRE | | | | |
| TC04001662.hg.1 | NM_000508 | FGA | fibrinogen alpha chain | -5.02 | 1.53E-05 | | | | | |
| TC03001692.hg.1 | NM_000187 | HGD | homogentisate 1,2-dioxygenase | -4.87 | 1.02E-05 | CRE | | | | |
| TC20000876.hg.1 | NM_003064 | SLPI | secretory leukocyte peptidase inhibitor | -4.76 | 0.000987 | CRE | | | | |
| TC03001719.hg.1 | NM_033049 | MUC13 | mucin 13, cell surface associated | -4.57 | 4.65E-06 | CRE | | | | |
| TC12000687.hg.1 | NM_006183 | NTS | neurotensin | -4.35 | 2.43E-06 | CRE | | | | |
| TC06001015.hg.1 | NM_018945 | PDE7B | phosphodiesterase 7B | -4.31 | 1.53E-05 | CRE | Yes | Yes | Yes | Yes |
| TC02002733.hg.1 | NM_001136574 | LANCL1 | LanC lantibiotic synthetase component C-like 1 (bacterial) | -4.29 | 6.14E-06 | | Yes | Yes | Yes | Yes |
| TC12000227.hg.1 | NM_000921 | PDE3A | phosphodiesterase 3A, cGMP-inhibited | -4.29 | 4.30E-06 | CRE | Yes | Yes | Yes | Yes |
| TC01003555.hg.1 | ENST0000042260 | PTP4A1P7 | protein tyrosine phosphatase type IVA, member 1 pseudogene 7 | -4.25 | 0.0001292 | | | | | |
| TC06002301.hg.1 | ENST0000054586 | SDIM1 | stress responsive DNAJB4 interacting membrane protein 1 | -4.16 | 0.0020345 | | | | | |
| TC12003227.hg.1 | NM_002281 | KRT81 | keratin 81 | -4.11 | 4.63E-06 | | | | | |
| TC08001690.hg.1 | NM_001080431 | SLC45A4 | solute carrier family 45, member 4 | -4.09 | 2.33E-05 | | Yes | Yes | Yes | Yes |
| TC01000733.hg.1 | NM_001037339 | PDE4B | phosphodiesterase 4B, cAMP-specific | -4.02 | 3.94E-06 | CRE | Yes | Yes | | |
| TC05001389.hg.1 | NM_001104631 | PDE4D | phosphodiesterase 4D, cAMP-specific | -3.98 | 1.49E-06 | CRE | Yes | Yes | Yes | Yes |
| TC02000281.hg.1 | NM_001430 | EPAS1 | endothelial PAS domain protein 1 | -3.92 | 3.94E-06 | CRE | | | Yes | Yes |
| TC07001512.hg.1 | ENST0000045042 | LOC101926943 | uncharacterized LOC101926943 | -3.92 | 8.64E-06 | | | | | |
| TC07003364.hg.1 | NM_000940 | PON3 | paraoxonase 3 | -3.82 | 7.89E-06 | CRE | Yes | Yes | Yes | Yes |
| TC16000501.hg.1 | NM_001142302 | CCDC113 | coiled-coil domain containing 113 | -3.61 | 7.12E-06 | CRE | Yes | Yes | Yes | Yes |
| TC19001657.hg.1 | NR_024258 | SNAR-E | small ILF3/NF90-associated RNA E | -3.56 | 0.0025273 | | | | | |
| TC16001140.hg.1 | NM_018110 | DOK4 | docking protein 4 | -3.55 | 1.02E-05 | CRE | Yes | Yes | Yes | |
| TC10000042.hg.1 | OTTHUMT000000 | RP11-499O7.7 | novel transcript, antisense to AKR1C2 | -3.55 | 0.0014298 | | | | | |
| TC07000276.hg.1 | NM_001003941 | OGDH | oxoglutarate (alpha-ketoglutarate) dehydrogenase (lipoamide) | -3.53 | 4.30E-06 | CRE | | | | |

| | | | | | | | | | | |
|---|---|---|---|---|---|---|---|---|---|---|
| TC02000219.hg.1 | NM_000627 | LTBP1 | latent transforming growth factor beta binding protein 1 | -3.49 | 9.60E-06 | CRE | Yes | Yes | Yes | Yes |
| TC14002318.hg.1 | NM_015473 | HEATR5A | HEAT repeat containing 5A | -3.47 | 8.72E-06 | | Yes | | Yes | |
| TC20000055.hg.1 | NR_028370 | PCNA-AS1 | PCNA antisense RNA 1 | -3.41 | 0.0001333 | | | | | |
| TC09000593.hg.1 | NM_002581 | PAPPA | pregnancy-associated plasma protein A, pappalysin 1 | -3.37 | 1.72E-05 | CRE | | | | |
| TC10002952.hg.1 | NM_001204300 | ARHGAP19 | Rho GTPase activating protein 19 | -3.29 | 1.39E-05 | | | | | |
| TC06001156.hg.1 | NM_021977 | SLC22A3 | solute carrier family 22 (organic cation transporter), member 3 | -3.18 | 2.19E-05 | CRE | Yes | Yes | Yes | Yes |
| TC19000321.hg.1 | NM_016270 | KLF2 | Kruppel-like factor 2 | -3.17 | 0.0005687 | CRE | | | | |
| TC0X000727.hg.1 | NM_005342 | HMGB3 | high mobility group box 3 | -3.13 | 5.40E-05 | CRE | | | | |
| TC16000497.hg.1 | AF452719 | LOC388282 | uncharacterized LOC388282 | -3.13 | 0.0011586 | | Yes | Yes | Yes | Yes |
| TC01004027.hg.1 | NM_001821 | CHML | choroideremia-like (Rab escort protein 2) | -3.12 | 0.0002592 | | | | | |
| TC10000449.hg.1 | NM_019058 | DDIT4 | DNA-damage-inducible transcript 4 | -3.10 | 3.36E-05 | CRE | Yes | Yes | Yes | Yes |
| TC11000464.hg.1 | NM_178570 | RTN4RL2 | reticulon 4 receptor-like 2 | -3.04 | 3.07E-05 | CRE | Yes | Yes | Yes | |
| TC05000097.hg.1 | ENST0000051709 | RNU6-1003P | RNA, U6 small nuclear 1003, pseudogene | -3.03 | 0.002992 | | | | | |
| TC12000311.hg.1 | NM_001256063 | CNTN1 | contactin 1 | -2.95 | 5.90E-06 | CRE | | | | |
| TC02000147.hg.1 | ENST0000036300 | RNU6-942P | RNA, U6 small nuclear 942, pseudogene | -2.94 | 0.0027632 | | | | | |
| TC06000697.hg.1 | NM_003463 | PTP4A1 | protein tyrosine phosphatase type IVA, member 1 | -2.92 | 1.27E-05 | CRE | | | | |
| TC03001512.hg.1 | NR_026582 | ID2B | inhibitor of DNA binding 2B, dominant negative helix-loop-helix proteir | -2.89 | 0.0025693 | | | | | |
| TC11001509.hg.1 | NM_018490 | LGR4 | leucine-rich repeat containing G protein-coupled receptor 4 | -2.83 | 1.52E-05 | CRE | | | | |
| TC08001014.hg.1 | NM_004467 | FGL1 | fibrinogen-like 1 | -2.82 | 3.82E-05 | CRE | | | | |
| TC20000398.hg.1 | ENST0000041145 | LINC01273 | long intergenic non-protein coding RNA 1273 | -2.79 | 0.0015591 | | Yes | Yes | Yes | Yes |
| TC11001414.hg.1 | NM_016422 | RNF141 | ring finger protein 141 | -2.74 | 0.0004913 | CRE | Yes | | Yes | |
| TC22000627.hg.1 | NM_001079539 | XBP1 | X-box binding protein 1 | -2.73 | 0.0001241 | CRE | Yes | Yes | | |
| TC14000936.hg.1 | NM_001126105 | SLC7A7 | solute carrier family 7 (amino acid transporter light chain, y+L system), r | -2.72 | 9.39E-05 | CRE | | | | |
| TC6_apd_hap1000 | NM_006120 | HLA-DMA | major histocompatibility complex, class II, DM alpha | -2.71 | 0.0003644 | | Yes | Yes | | |
| TC13000295.hg.1 | NM_001160706 | SCEL | sciellin | -2.68 | 0.0002176 | CRE | | | | |
| TC12001363.hg.1 | NM_144973 | DENND5B | DENN/MADD domain containing 5B | -2.68 | 1.45E-05 | CRE | Yes | Yes | Yes | Yes |
| TC0X000901.hg.1 | NM_001079858 | GPR64 | G protein-coupled receptor 64 | -2.67 | 9.10E-05 | CRE | | | | |
| TC01001976.hg.1 | NM_003686 | EXO1 | exonuclease 1 | -2.66 | 1.27E-05 | CRE | Yes | Yes | Yes | Yes |
| TC19000688.hg.1 | NR_004435 | SNAR-A1 | small ILF3/NF90-associated RNA A1 | -2.63 | 0.0002188 | | | | | |
| TC19000691.hg.1 | NR_004436 | SNAR-A2 | small ILF3/NF90-associated RNA A2 | -2.63 | 0.0002188 | | | | | |
| TC19000689.hg.1 | NR_024214 | SNAR-A3 | small ILF3/NF90-associated RNA A3 | -2.63 | 0.0002188 | | | | | |
| TC19001731.hg.1 | NR_024242 | SNAR-A14 | small ILF3/NF90-associated RNA A14 | -2.63 | 0.0002188 | | | | | |
| TC10000967.hg.1 | NM_004508 | IDI1 | isopentenyl-diphosphate delta isomerase 1 | -2.62 | 0.0001696 | CRE | Yes | Yes | Yes | Yes |
| TC03000892.hg.1 | NM_001122752 | SERPINI1 | serpin peptidase inhibitor, clade I (neuroserpin), member 1 | -2.61 | 0.0001099 | CRE | | | Yes | Yes |
| TC06001271.hg.1 | NM_017770 | ELOVL2 | ELOVL fatty acid elongase 2 | -2.61 | 0.0010171 | CRE | Yes | | Yes | |
| TC09001001.hg.1 | NM_001199987 | NDUFB6 | NADH dehydrogenase (ubiquinone) 1 beta subcomplex, 6, 17kDa | -2.60 | 0.0001066 | CRE | | | Yes | Yes |
| TC20000588.hg.1 | NM_005116 | SLC23A2 | solute carrier family 23 (ascorbic acid transporter), member 2 | -2.60 | 0.0001193 | | | | | |
| TC01001736.hg.1 | NM_000574 | CD55 | CD55 molecule, decay accelerating factor for complement (Cromer bloc | -2.58 | 2.02E-05 | CRE | Yes | | Yes | |
| TC01002347.hg.1 | NM_002167 | ID3 | inhibitor of DNA binding 3, dominant negative helix-loop-helix protein | -2.58 | 0.0016626 | CRE | | | | |
| TC06001298.hg.1 | NM_001105566 | KIF13A | kinesin family member 13A | -2.55 | 2.36E-05 | CRE | Yes | Yes | | |
| TC08000132.hg.1 | ENST0000052064 | LOC101930275 | uncharacterized LOC101930275 | -2.55 | 0.002822 | | | | | |
| TC11001999.hg.1 | NM_030930 | UNC93B1 | unc-93 homolog B1 (C. elegans) | -2.53 | 7.58E-05 | CRE | | | | |
| TC17001698.hg.1 | NM_001243877 | TOB1 | transducer of ERBB2, 1 | -2.53 | 4.51E-05 | CRE | Yes | Yes | Yes | Yes |
| TC6_mann_hap40( | NM_002118 | HLA-DMB | major histocompatibility complex, class II, DM beta | -2.51 | 9.39E-05 | CRE | | | | |
| TC02001719.hg.1 | ENST0000054262 | KRT18P52 | keratin 18 pseudogene 52 | -2.50 | 6.92E-05 | | | | | |
| TC07003361.hg.1 | NM_000786 | CYP51A1 | cytochrome P450, family 51, subfamily A, polypeptide 1 | -2.48 | 5.64E-05 | CRE | Yes | | Yes | |
| TC01004000.hg.1 | ENST0000051624 | RNU5E-2P | RNA, U5E small nuclear 2, pseudogene | -2.45 | 0.0112626 | | | | | |
| TC18000272.hg.1 | NM_005131 | THOC1 | THO complex 1 | -2.44 | 4.22E-05 | CRE | Yes | Yes | Yes | Yes |
| TC19000686.hg.1 | NR_004437 | SNAR-A12 | small ILF3/NF90-associated RNA A12 | -2.44 | 0.000181 | | | | | |
| TC19000693.hg.1 | NR_024216 | SNAR-A13 | small ILF3/NF90-associated RNA A13 | -2.44 | 0.000181 | | | | | |
| TC07000023.hg.1 | NM_002360 | MAFK | v-maf avian musculoaponeurotic fibrosarcoma oncogene homolog K | -2.43 | 0.0003625 | CRE | Yes | | | |

| | | | | | | | | | | |
|---|---|---|---|---|---|---|---|---|---|---|
| TC19000055.hg.1 | NM_015675 | GADD45B | growth arrest and DNA-damage-inducible, beta | -2.42 | 0.0040271 | CRE | Yes | | Yes | Yes |
| TC16002038.hg.1 | NM_001014449 | DDX19B | DEAD (Asp-Glu-Ala-Asp) box polypeptide 19B | -2.40 | 3.36E-05 | CRE | | | Yes | Yes |
| TC02002676.hg.1 | NM_001044385 | TMEM237 | transmembrane protein 237 | -2.40 | 3.07E-05 | CRE | Yes | Yes | Yes | Yes |
| TC15000124.hg.1 | NM_000810 | GABRA5 | gamma-aminobutyric acid (GABA) A receptor, alpha 5 | -2.39 | 9.35E-05 | CRE | | | Yes | Yes |
| TC07001828.hg.1 | NM_005302 | GPR37 | G protein-coupled receptor 37 (endothelin receptor type B-like) | -2.38 | 0.0016194 | CRE | | | | |
| TC05000096.hg.1 | NM_006317 | BASP1 | brain abundant, membrane attached signal protein 1 | -2.35 | 0.0004851 | CRE | | | Yes | Yes |
| TC01003968.hg.1 | NM_022051 | EGLN1 | egl-9 family hypoxia-inducible factor 1 | -2.35 | 5.36E-05 | CRE | | | | |
| TC18000524.hg.1 | NM_001143829 | CCDC68 | coiled-coil domain containing 68 | -2.35 | 0.0004115 | | Yes | Yes | Yes | Yes |
| TC09001009.hg.1 | NM_001172415 | BAG1 | BCL2-associated athanogene | -2.34 | 7.03E-05 | | Yes | Yes | Yes | |
| TC19000014.hg.1 | NM_173481 | MISP | mitotic spindle positioning | -2.33 | 0.0017787 | CRE | | | Yes | Yes |
| TC05001184.hg.1 | NM_012334 | MYO10 | myosin X | -2.33 | 0.0001333 | CRE | | | | |
| TC08001013.hg.1 | NM_001001924 | MTUS1 | microtubule associated tumor suppressor 1 | -2.32 | 0.0001684 | CRE | Yes | Yes | Yes | Yes |
| TC03000051.hg.1 | NM_001018115 | FANCD2 | Fanconi anemia, complementation group D2 | -2.30 | 0.0001334 | CRE | Yes | Yes | Yes | Yes |
| TC12001170.hg.1 | NM_006931 | SLC2A3 | solute carrier family 2 (facilitated glucose transporter), member 3 | -2.30 | 7.36E-05 | | | | | |
| TC06000062.hg.1 | NM_001718 | BMP6 | bone morphogenetic protein 6 | -2.28 | 0.0001099 | | Yes | Yes | | |
| TC08001022.hg.1 | NM_001130518 | CSGALNACT1 | chondroitin sulfate N-acetylgalactosaminyltransferase 1 | -2.27 | 3.45E-05 | | Yes | Yes | Yes | Yes |
| TC17001296.hg.1 | NM_000638 | VTN | vitronectin | -2.27 | 0.000263 | CRE | Yes | Yes | Yes | Yes |
| TC20000082.hg.1 | NM_003081 | SNAP25 | synaptosomal-associated protein, 25kDa | -2.26 | 0.0001191 | CRE | Yes | Yes | Yes | Yes |
| TC01003341.hg.1 | NM_005920 | MEF2D | myocyte enhancer factor 2D | -2.25 | 6.36E-05 | CRE | Yes | Yes | | |
| TC07000569.hg.1 | NM_016116 | ASB4 | ankyrin repeat and SOCS box containing 4 | -2.24 | 0.0003956 | CRE | | | | |
| TC0X000088.hg.1 | NM_001037343 | CDKL5 | cyclin-dependent kinase-like 5 | -2.24 | 0.0001839 | | Yes | Yes | Yes | Yes |
| TC13000871.hg.1 | NM_003749 | IRS2 | insulin receptor substrate 2 | -2.23 | 0.0001241 | | | | | |
| TC15000060.hg.1 | NR_003329 | SNORD116-14 | small nucleolar RNA, C/D box 116-14 | -2.23 | 0.0003298 | | | | | |
| TC08000127.hg.1 | NM_001008539 | SLC7A2 | solute carrier family 7 (cationic amino acid transporter, y+ system), men | -2.22 | 0.0002613 | | Yes | Yes | | |
| TC17000495.hg.1 | NM_001254 | CDC6 | cell division cycle 6 | -2.20 | 0.0001105 | CRE | Yes | Yes | Yes | Yes |
| TC15000824.hg.1 | NM_001243137 | PDE8A | phosphodiesterase 8A | -2.19 | 5.96E-05 | CRE | Yes | | Yes | |
| TC01001781.hg.1 | NM_014053 | FLVCR1 | feline leukemia virus subgroup C cellular receptor 1 | -2.18 | 0.0002786 | CRE | Yes | | Yes | |
| TC04001219.hg.1 | NM_181806 | AASDH | aminoadipate-semialdehyde dehydrogenase | -2.18 | 0.0008147 | CRE | Yes | Yes | Yes | Yes |
| TC17000583.hg.1 | NR_024434 | MAP3K14-AS1 | MAP3K14 antisense RNA 1 | -2.18 | 0.0011347 | | Yes | Yes | Yes | Yes |
| TC09001325.hg.1 | NM_005384 | NFIL3 | nuclear factor, interleukin 3 regulated | -2.18 | 0.0008391 | CRE | Yes | Yes | Yes | Yes |
| TC15000052.hg.1 | NR_003317 | SNORD116-2 | small nucleolar RNA, C/D box 116-2 | -2.17 | 0.0007056 | | | | | |
| TC07000692.hg.1 | NM_181581 | DUS4L | dihydrouridine synthase 4-like (S. cerevisiae) | -2.17 | 0.0001099 | CRE | Yes | Yes | Yes | Yes |
| TC19001139.hg.1 | NM_024690 | MUC16 | mucin 16, cell surface associated | -2.17 | 0.0001502 | | | | | |
| TC01002769.hg.1 | NM_017768 | LRRC40 | leucine rich repeat containing 40 | -2.16 | 7.56E-05 | CRE | | | Yes | Yes |
| TC15000304.hg.1 | NM_007236 | CHP1 | calcineurin-like EF-hand protein 1 | -2.15 | 3.82E-05 | CRE | Yes | Yes | Yes | Yes |
| TC19000350.hg.1 | NM_001025604 | ARRDC2 | arrestin domain containing 2 | -2.14 | 0.0003608 | CRE | Yes | Yes | Yes | Yes |
| TC19001278.hg.1 | NM_001080421 | UNC13A | unc-13 homolog A (C. elegans) | -2.12 | 3.56E-05 | | Yes | Yes | Yes | Yes |
| TC02001528.hg.1 | BC113076 | TMSB4XP2 | thymosin beta 4, X-linked pseudogene 2 | -2.12 | 0.0014133 | CRE | | | | |
| TC01004026.hg.1 | NM_014322 | OPN3 | opsin 3 | -2.10 | 0.0002412 | CRE | | | | |
| TC10001320.hg.1 | NM_001242359 | RHOBTB1 | Rho-related BTB domain containing 1 | -2.10 | 0.0002286 | CRE | Yes | Yes | Yes | Yes |
| TC04000416.hg.1 | NM_015393 | PARM1 | prostate androgen-regulated mucin-like protein 1 | -2.10 | 0.0002057 | CRE | | | | |
| TC08001099.hg.1 | NM_001394 | DUSP4 | dual specificity phosphatase 4 | -2.10 | 0.0001431 | CRE | Yes | Yes | | |
| TC6_mcf_hap5000 | NR_037177 | LOC100294145 | uncharacterized LOC100294145 | -2.10 | 0.0019611 | | | | | |
| TC09001166.hg.1 | ENST0000045957 | MIR1299 | microRNA 1299 | -2.09 | 0.000834 | | | | | |
| TC04001765.hg.1 | NM_005429 | VEGFC | vascular endothelial growth factor C | -2.09 | 0.0005418 | CRE | | | Yes | Yes |
| TC15001084.hg.1 | NM_000814 | GABRB3 | gamma-aminobutyric acid (GABA) A receptor, beta 3 | -2.08 | 0.000217 | | Yes | Yes | Yes | Yes |
| TC06001658.hg.1 | NM_001252294 | SPDEF | SAM pointed domain containing ETS transcription factor | -2.08 | 0.0001315 | CRE | | | | |
| TC10000098.hg.1 | NM_018518 | MCM10 | minichromosome maintenance complex component 10 | -2.07 | 0.0001574 | CRE | Yes | Yes | Yes | Yes |
| TC20000049.hg.1 | NR_033917 | LINC01433 | long intergenic non-protein coding RNA 1433 | -2.06 | 0.0062753 | | Yes | Yes | Yes | Yes |
| TC14000391.hg.1 | NM_021979 | HSPA2 | heat shock 70kDa protein 2 | -2.06 | 0.0022013 | | Yes | Yes | Yes | |
| TC03000351.hg.1 | NM_000720 | CACNA1D | calcium channel, voltage-dependent, L type, alpha 1D subunit | -2.06 | 0.0001191 | CRE | Yes | | Yes | |

| | | | | | | | | | | |
|---|---|---|---|---|---|---|---|---|---|---|
| TC14000202.hg.1 | NM_001201573 | NUBPL | nucleotide binding protein-like | -2.05 | 0.0002188 | CRE | Yes | Yes | | |
| TC0X001155.hg.1 | NR_001564 | XIST | X inactive specific transcript (non-protein coding) | -2.04 | 0.0040543 | | | | | |
| TC0X002360.hg.1 | NR_038988 | LINC00630 | long intergenic non-protein coding RNA 630 | -2.04 | 0.0030757 | | Yes | Yes | | |
| TC15000059.hg.1 | NR_003328 | SNORD116-13 | small nucleolar RNA, C/D box 116-13 | -2.04 | 0.00493 | | | | | |
| TC11002035.hg.1 | OTTHUMT000002 | SHANK2 | SH3 and multiple ankyrin repeat domains 2 | -2.04 | 0.0254243 | | | | | |
| TC15000050.hg.1 | NR_003318 | SNORD116-3 | small nucleolar RNA, C/D box 116-3 | -2.04 | 0.0014403 | | | | | |
| TC15000055.hg.1 | NR_003324 | SNORD116-9 | small nucleolar RNA, C/D box 116-9 | -2.04 | 0.0014403 | | | | | |
| TC01001741.hg.1 | NM_002389 | CD46 | CD46 molecule, complement regulatory protein | -2.04 | 0.0001585 | CRE | | | | |
| TC15000305.hg.1 | NR_026757 | OIP5-AS1 | OIP5 antisense RNA 1 | -2.04 | 0.0003058 | | Yes | Yes | | |
| TC19000020.hg.1 | NM_001928 | CFD | complement factor D (adipsin) | -2.03 | 0.0006671 | CRE | Yes | Yes | Yes | Yes |
| TC02001627.hg.1 | NM_021925 | C2orf43 | chromosome 2 open reading frame 43 | -2.02 | 0.0004307 | CRE | | | | |
| TC02000612.hg.1 | OTTHUMT000003 | AC092168.2 | novel transcript | -2.02 | 0.022266 | | | | | |
| TC06000012.hg.1 | NM_001453 | FOXC1 | forkhead box C1 | -2.01 | 0.0001502 | CRE | Yes | | Yes | Yes |
| TC17000580.hg.1 | NM_006460 | HEXIM1 | hexamethylene bis-acetamide inducible 1 | -2.00 | 0.0004628 | CRE | | | | |
| TC17001465.hg.1 | NM_032865 | TNS4 | tensin 4 | -2.00 | 0.0008728 | CRE | Yes | Yes | Yes | Yes |
| TC03001629.hg.1 | NM_001777 | CD47 | CD47 molecule | 2.00 | 0.0004895 | CRE | Yes | Yes | Yes | Yes |
| TC07001080.hg.1 | NM_182491 | ZFAND2A | zinc finger, AN1-type domain 2A | 2.01 | 0.0001574 | CRE | | | | |
| TC01000745.hg.1 | NM_001199741 | GADD45A | growth arrest and DNA-damage-inducible, alpha | 2.01 | 0.0001315 | | | | | |
| TC02001696.hg.1 | NR_028308 | BRE-AS1 | BRE antisense RNA 1 | 2.01 | 0.0386721 | | | | | |
| TC05001385.hg.1 | NM_001252226 | PLK2 | polo-like kinase 2 | 2.02 | 0.0001574 | CRE | Yes | Yes | Yes | Yes |
| TC02002707.hg.1 | ENST0000042877 | KLF7-IT1 | KLF7 intronic transcript 1 (non-protein coding) | 2.02 | 0.0033904 | | | | | |
| TC05000083.hg.1 | NM_007118 | TRIO | trio Rho guanine nucleotide exchange factor | 2.02 | 0.0002624 | CRE | Yes | Yes | | |
| TC06000721.hg.1 | OTTHUMT000000 | RP3-331H24.4 | putative novel transcript | 2.02 | 0.009203 | | | | | |
| TC12000611.hg.1 | NM_000239 | LYZ | lysozyme | 2.03 | 0.0022135 | | | | | |
| TC06004141.hg.1 | NM_000636 | SOD2 | superoxide dismutase 2, mitochondrial | 2.03 | 0.0014942 | CRE | Yes | Yes | Yes | Yes |
| TC07001293.hg.1 | NM_003014 | SFRP4 | secreted frizzled-related protein 4 | 2.03 | 0.000236 | CRE | | | | |
| TC12001178.hg.1 | NM_014358 | CLEC4E | C-type lectin domain family 4, member E | 2.03 | 0.0023127 | CRE | | | | |
| TC02002419.hg.1 | NM_001254738 | RND3 | Rho family GTPase 3 | 2.04 | 0.0001439 | CRE | Yes | Yes | Yes | Yes |
| TC05001861.hg.1 | NM_000591 | CD14 | CD14 molecule | 2.04 | 0.0007612 | | | | | |
| TC03001887.hg.1 | NM_001184723 | TM4SF18 | transmembrane 4 L six family member 18 | 2.04 | 0.0003032 | CRE | | | | |
| TC03001651.hg.1 | NM_199511 | CCDC80 | coiled-coil domain containing 80 | 2.04 | 0.0002124 | | | | | |
| TC15000089.hg.1 | NR_003309 | SNORD115-17 | small nucleolar RNA, C/D box 115-17 | 2.05 | 0.0460187 | | | | | |
| TC15000090.hg.1 | NR_003310 | SNORD115-18 | small nucleolar RNA, C/D box 115-18 | 2.05 | 0.0460187 | | | | | |
| TC15000091.hg.1 | NR_003311 | SNORD115-19 | small nucleolar RNA, C/D box 115-19 | 2.05 | 0.0460187 | | | | | |
| TC04001283.hg.1 | NM_002994 | CXCL5 | chemokine (C-X-C motif) ligand 5 | 2.06 | 0.0004029 | CRE | Yes | Yes | Yes | Yes |
| TC02000275.hg.1 | NM_000341 | SLC3A1 | solute carrier family 3 (amino acid transporter heavy chain), member 1 | 2.06 | 0.0001138 | | | | | |
| TC15000622.hg.1 | NM_001145102 | SMAD3 | SMAD family member 3 | 2.06 | 0.0001433 | CRE | Yes | Yes | Yes | Yes |
| TC03001869.hg.1 | NM_021105 | PLSCR1 | phospholipid scramblase 1 | 2.06 | 0.0001001 | | | | | |
| TC01003007.hg.1 | NM_001010922 | BCL2L15 | BCL2-like 15 | 2.06 | 0.0008291 | | | | | |
| TC21000200.hg.1 | AL355711 | ABCG1 | ATP-binding cassette, sub-family G (WHITE), member 1 | 2.08 | 0.0013618 | | Yes | Yes | | |
| TC02002481.hg.1 | NM_022168 | IFIH1 | interferon induced with helicase C domain 1 | 2.08 | 0.0001992 | CRE | Yes | | Yes | |
| TC07000625.hg.1 | NM_006076 | AGFG2 | ArfGAP with FG repeats 2 | 2.09 | 9.35E-05 | CRE | Yes | Yes | Yes | Yes |
| TC02002827.hg.1 | NM_014689 | DOCK10 | dedicator of cytokinesis 10 | 2.09 | 0.0001839 | CRE | | | | |
| TC12001783.hg.1 | NM_001146335 | SLC6A15 | solute carrier family 6 (neutral amino acid transporter), member 15 | 2.09 | 0.0002488 | CRE | | | Yes | Yes |
| TC05000243.hg.1 | NM_005921 | MAP3K1 | mitogen-activated protein kinase kinase kinase 1, E3 ubiquitin protein li | 2.11 | 0.0002188 | | Yes | | Yes | |
| TC03001955.hg.1 | ENST0000045881 | RNU7-136P | RNA, U7 small nuclear 136 pseudogene | 2.11 | 0.0268382 | | | | | |
| TC11000252.hg.1 | NM_001111018 | NAV2 | neuron navigator 2 | 2.11 | 0.0001907 | CRE | Yes | | Yes | Yes |
| TC02001140.hg.1 | ENST0000043393 | LOC101927482 | uncharacterized LOC101927482 | 2.11 | 0.0012974 | | | | | |
| TC16000879.hg.1 | NM_001099455 | CPPED1 | calcineurin-like phosphoesterase domain containing 1 | 2.14 | 0.0003632 | CRE | | | | |
| TC01002903.hg.1 | NM_000110 | DPYD | dihydropyrimidine dehydrogenase | 2.14 | 9.43E-05 | | | | Yes | Yes |
| TC15001719.hg.1 | NM_001114735 | BCL2A1 | BCL2-related protein A1 | 2.14 | 0.0110898 | | | | | |

| | | | | | | | | | | |
|---|---|---|---|---|---|---|---|---|---|---|
| TC11001107.hg.1 | NM_003105 | SORL1 | sortilin-related receptor, L(DLR class) A repeats containing | 2.14 | 0.0001439 | CRE | Yes | Yes | | |
| TC06001128.hg.1 | NM_001178088 | SYNJ2 | synaptojanin 2 | 2.14 | 6.68E-05 | CRE | Yes | Yes | Yes | Yes |
| TC17001924.hg.1 | NM_005567 | LGALS3BP | lectin, galactoside-binding, soluble, 3 binding protein | 2.15 | 0.0017242 | | | | Yes | Yes |
| TC03000498.hg.1 | ENST0000047375 | LINC00973 | long intergenic non-protein coding RNA 973 | 2.17 | 0.0021865 | | | | | |
| TC06001035.hg.1 | NM_016217 | HECA | headcase homolog (Drosophila) | 2.17 | 0.0007644 | CRE | | | | |
| TC04000912.hg.1 | NM_003265 | TLR3 | toll-like receptor 3 | 2.18 | 0.0001099 | | | | | |
| TC01003267.hg.1 | NM_002960 | S100A3 | S100 calcium binding protein A3 | 2.18 | 0.0001463 | CRE | Yes | Yes | | |
| TC02002823.hg.1 | NM_001136528 | SERPINE2 | serpin peptidase inhibitor, clade E (nexin, plasminogen activator inhibit | 2.18 | 8.68E-05 | CRE | Yes | | Yes | |
| TC16000921.hg.1 | NM_016235 | GPRC5B | G protein-coupled receptor, class C, group 5, member B | 2.18 | 0.0002911 | CRE | Yes | Yes | Yes | Yes |
| TC14001238.hg.1 | NM_016445 | PLEK2 | pleckstrin 2 | 2.18 | 0.0005332 | CRE | Yes | Yes | | |
| TC08001253.hg.1 | NM_004056 | CA8 | carbonic anhydrase VIII | 2.20 | 0.0001433 | | | | | |
| TC01003847.hg.1 | NM_007207 | DUSP10 | dual specificity phosphatase 10 | 2.21 | 0.0001464 | CRE | Yes | Yes | Yes | Yes |
| TC06002242.hg.1 | NM_012419 | RGS17 | regulator of G-protein signaling 17 | 2.21 | 0.0007735 | CRE | Yes | Yes | Yes | Yes |
| TC16000214.hg.1 | NM_001105248 | TMC5 | transmembrane channel-like 5 | 2.22 | 0.0002072 | | Yes | Yes | Yes | Yes |
| TC12001705.hg.1 | NM_001005502 | CPM | carboxypeptidase M | 2.22 | 0.0004301 | CRE | Yes | Yes | Yes | Yes |
| TC15000752.hg.1 | NM_018689 | CEMIP | cell migration inducing protein, hyaluronan binding | 2.23 | 0.0050449 | | Yes | Yes | Yes | Yes |
| TC20000913.hg.1 | NM_001161841 | SULF2 | sulfatase 2 | 2.25 | 0.0001696 | CRE | Yes | Yes | Yes | Yes |
| TC10000420.hg.1 | NM_012339 | TSPAN15 | tetraspanin 15 | 2.25 | 4.89E-05 | | | | | |
| TC02002891.hg.1 | NM_005737 | ARL4C | ADP-ribosylation factor-like 4C | 2.27 | 0.0001574 | CRE | Yes | Yes | Yes | Yes |
| TC09001276.hg.1 | NM_152573 | RASEF | RAS and EF-hand domain containing | 2.27 | 4.23E-05 | CRE | | | | |
| TC01003784.hg.1 | NM_000228 | LAMB3 | laminin, beta 3 | 2.31 | 0.0001241 | CRE | Yes | Yes | | |
| TC01002654.hg.1 | NM_002867 | RAB3B | RAB3B, member RAS oncogene family | 2.31 | 9.19E-05 | CRE | | | | |
| TC01000288.hg.1 | NM_004442 | EPHB2 | EPH receptor B2 | 2.31 | 0.000182 | | Yes | Yes | Yes | Yes |
| TC01000814.hg.1 | NM_145172 | WDR63 | WD repeat domain 63 | 2.33 | 0.0006083 | CRE | Yes | Yes | | |
| TC0X000085.hg.1 | NR_039925 | MIR4768 | microRNA 4768 | 2.34 | 0.0019008 | | | | | |
| TC04001830.hg.1 | NM_173553 | TRIML2 | tripartite motif family-like 2 | 2.35 | 0.0013707 | CRE | | | | |
| TC02001127.hg.1 | ENST0000051632 | RNU6-1045P | RNA, U6 small nuclear 1045, pseudogene | 2.36 | 0.0391042 | | | | | |
| TC02002627.hg.1 | NM_004657 | SDPR | serum deprivation response | 2.36 | 0.019058 | CRE | | | | |
| TC14001253.hg.1 | NM_001244698 | ZFP36L1 | ZFP36 ring finger protein-like 1 | 2.38 | 7.90E-05 | CRE | Yes | | Yes | |
| TC07001611.hg.1 | NM_006528 | TFPI2 | tissue factor pathway inhibitor 2 | 2.39 | 8.68E-05 | CRE | | | | |
| TC09000560.hg.1 | NM_003358 | UGCG | UDP-glucose ceramide glucosyltransferase | 2.42 | 2.87E-05 | | | | | |
| TC01001930.hg.1 | NM_173508 | SLC35F3 | solute carrier family 35, member F3 | 2.43 | 0.0002114 | CRE | | | Yes | Yes |
| TC14001474.hg.1 | NM_024734 | CLMN | calmin (calponin-like, transmembrane) | 2.50 | 0.0001307 | CRE | Yes | Yes | Yes | Yes |
| TC03000831.hg.1 | NM_033050 | SUCNR1 | succinate receptor 1 | 2.55 | 0.0002939 | CRE | | | | |
| TC01006361.hg.1 | NM_004120 | GBP2 | guanylate binding protein 2, interferon-inducible | 2.56 | 1.89E-05 | CRE | | | | |
| TC19000584.hg.1 | NM_002483 | CEACAM6 | carcinoembryonic antigen-related cell adhesion molecule 6 (non-specifi | 2.56 | 0.0004157 | | | | | |
| TC17001378.hg.1 | NM_002985 | CCL5 | chemokine (C-C motif) ligand 5 | 2.57 | 0.000407 | CRE | | | | |
| TC02000762.hg.1 | NM_002193 | INHBB | inhibin, beta B | 2.57 | 0.0004976 | CRE | Yes | | Yes | |
| TC04000408.hg.1 | NM_000584 | CXCL8 | chemokine (C-X-C motif) ligand 8 | 2.57 | 0.0002525 | CRE | | | | |
| TC09001618.hg.1 | NM_001035254 | FAM102A | family with sequence similarity 102, member A | 2.58 | 6.92E-05 | | Yes | Yes | Yes | Yes |
| TC01002163.hg.1 | NM_001561 | TNFRSF9 | tumor necrosis factor receptor superfamily, member 9 | 2.59 | 0.0002132 | CRE | | | | |
| TC06002152.hg.1 | NM_000416 | IFNGR1 | interferon gamma receptor 1 | 2.59 | 8.64E-06 | CRE | | | | |
| TC19001747.hg.1 | NM_138411 | FAM71E1 | family with sequence similarity 71, member E1 | 2.61 | 0.0004213 | | | | | |
| TC03002096.hg.1 | NM_001130845 | BCL6 | B-cell CLL/lymphoma 6 | 2.62 | 8.68E-05 | CRE | | | | |
| TC17000861.hg.1 | NM_000213 | ITGB4 | integrin, beta 4 | 2.63 | 2.02E-05 | CRE | Yes | | Yes | Yes |
| TC17001250.hg.1 | NM_000691 | ALDH3A1 | aldehyde dehydrogenase 3 family, member A1 | 2.63 | 0.0002862 | CRE | Yes | Yes | Yes | Yes |
| TC08001440.hg.1 | NM_001135733 | TP53INP1 | tumor protein p53 inducible nuclear protein 1 | 2.64 | 1.74E-05 | CRE | Yes | Yes | Yes | Yes |
| TC15001641.hg.1 | NM_001142617 | STRA6 | stimulated by retinoic acid 6 | 2.64 | 0.0001241 | | Yes | Yes | Yes | Yes |
| TC01002287.hg.1 | NM_007365 | PADI2 | peptidyl arginine deiminase, type II | 2.65 | 6.45E-05 | CRE | Yes | Yes | Yes | Yes |
| TC01000955.hg.1 | NM_000757 | CSF1 | colony stimulating factor 1 (macrophage) | 2.65 | 0.0001574 | | | | | |
| TC0X000879.hg.1 | NM_020665 | TMEM27 | transmembrane protein 27 | 2.72 | 0.0006713 | CRE | | | | |

| | | | | | | | | | | |
|---|---|---|---|---|---|---|---|---|---|---|
| TC21000480.hg.1 | NM_001098402 | ZBTB21 | zinc finger and BTB domain containing 21 | 2.72 | 2.33E-05 | | | | Yes | |
| TC19000174.hg.1 | NM_000201 | ICAM1 | intercellular adhesion molecule 1 | 2.74 | 0.0003678 | CRE | Yes | | Yes | |
| TC02002746.hg.1 | NM_015657 | ABCA12 | ATP-binding cassette, sub-family A (ABC1), member 12 | 2.76 | 7.29E-05 | CRE | | | | |
| TC05002043.hg.1 | NM_003062 | SLIT3 | slit homolog 3 (Drosophila) | 2.82 | 1.43E-05 | CRE | | | | |
| TC01000723.hg.1 | NM_001083592 | ROR1 | receptor tyrosine kinase-like orphan receptor 1 | 2.82 | 0.0001817 | CRE | | | | |
| TC04000411.hg.1 | NM_001511 | CXCL1 | chemokine (C-X-C motif) ligand 1 (melanoma growth stimulating activit | 2.98 | 0.0005801 | CRE | | | Yes | Yes |
| TC11000956.hg.1 | NM_001165 | BIRC3 | baculoviral IAP repeat containing 3 | 3.01 | 9.39E-05 | CRE | | | | |
| TC14002309.hg.1 | NM_001085 | SERPINA3 | serpin peptidase inhibitor, clade A (alpha-1 antiproteinase, antitrypsin), | 3.11 | 3.11E-05 | CRE | | | | |
| TC03000056.hg.1 | NM_001570 | IRAK2 | interleukin-1 receptor-associated kinase 2 | 3.15 | 1.52E-05 | CRE | | | | |
| TC01000309.hg.1 | NM_001195010 | GRHL3 | grainyhead-like 3 (Drosophila) | 3.18 | 5.85E-05 | | Yes | | Yes | |
| TC14000794.hg.1 | NM_006291 | TNFAIP2 | tumor necrosis factor, alpha-induced protein 2 | 3.30 | 3.45E-05 | CRE | | | | |
| TC06001027.hg.1 | NM_006290 | TNFAIP3 | tumor necrosis factor, alpha-induced protein 3 | 3.30 | 1.60E-05 | CRE | Yes | Yes | Yes | Yes |
| TC07000643.hg.1 | NM_000602 | SERPINE1 | serpin peptidase inhibitor, clade E (nexin, plasminogen activator inhibit | 3.41 | 4.22E-05 | CRE | Yes | Yes | Yes | Yes |
| TC12000656.hg.1 | NM_014903 | NAV3 | neuron navigator 3 | 3.52 | 3.94E-06 | CRE | | | | |
| TC05002146.hg.1 | NM_005110 | GFPT2 | glutamine-fructose-6-phosphate transaminase 2 | 3.54 | 3.94E-06 | | Yes | Yes | Yes | Yes |
| TC19001101.hg.1 | NM_001252 | CD70 | CD70 molecule | 3.97 | 0.0001839 | | | | Yes | Yes |
| TC20000718.hg.1 | NM_001898 | CST1 | cystatin SN | 4.14 | 7.49E-05 | | | | | |
| TC12000189.hg.1 | NM_001423 | EMP1 | epithelial membrane protein 1 | 4.38 | 2.14E-05 | CRE | | | | |
| TC10000475.hg.1 | NM_001145031 | PLAU | plasminogen activator, urokinase | 4.48 | 5.24E-06 | CRE | | | | |
| TC20000717.hg.1 | NM_001899 | CST4 | cystatin S | 4.57 | 7.36E-05 | | | | | |
| TC15000971.hg.1 | NM_000693 | ALDH1A3 | aldehyde dehydrogenase 1 family, member A3 | 4.95 | 1.88E-05 | CRE | Yes | Yes | Yes | Yes |
| TC19001644.hg.1 | BC062328 | IGFL2-AS1 | IGFL2 antisense RNA 1 | 4.98 | 8.68E-05 | | | | | |
| TC01000231.hg.1 | NM_016233 | PADI3 | peptidyl arginine deiminase, type III | 5.74 | 2.43E-06 | CRE | | | | |
| TC10001517.hg.1 | NM_014391 | ANKRD1 | ankyrin repeat domain 1 (cardiac muscle) | 6.03 | 1.52E-05 | | | | | |
| TC07003299.hg.1 | NM_000777 | CYP3A5 | cytochrome P450, family 3, subfamily A, polypeptide 5 | 6.52 | 6.94E-07 | | | | | |
| TC17000383.hg.1 | NM_002982 | CCL2 | chemokine (C-C motif) ligand 2 | 12.58 | 1.03E-05 | | | | | |
| TC11002228.hg.1 | NM_002423 | MMP7 | matrix metallopeptidase 7 (matrilysin, uterine) | 25.46 | 6.94E-07 | CRE | Yes | Yes | Yes | Yes |

**Table S2: Primer and sgRNA sequences used in this study. Sequences were presented in the 5' to 3' direction.**

**RT-PCR primers:**

| Gene | Forward Primer | Reverse Primer | Amplication (bp) |
|---|---|---|---|
| CRTC1 | TGTCTCTCTGACCCCCTTCCAATCC | GTCCGCGGGTGGTGAGAGGTA | 196 |
| CRTC2 | AGCCCCCTGAGTTTGCTCGC | TGGGGGTAACCGCTGGTCAGT | 159 |
| CRTC3 | TGACCAGCAGTCCATGAGGCCA | GGTCTTTGAACAGGCTGGTGCTGG | 176 |
| LINC00473 | AAACGCGAACGTGAGCCCCG | CGCCATGCTCTGGCGCAGTT | 134 |
| FOS | CACTCCAAGCGGAGACAG | AGGTCATCAGGGATCTTGCAG | 139 |
| NR4A2 | GCCGGAGAGGTCGTTTGCCC | AGGGTTCGCCTGGAACCTGGAA | 155 |
| INSL4 | GATGTGGTCCCCGATTTGGA | AGGTTGACACCATTTCTTTGGG | 123 |
| CPS1 | CTGATGCTGCCCACACAAAC | AGGGGAAGGATCGAGAAGCT | 175 |
| PDK4 | ACAGACAGGAAACCCAAGCC | GTTCAACTGTTGCCCGCATT | 248 |
| NR4A1 | GAGTCCCAGTGGCGGAGGCT | CAGGCTGCACCCTACCCGGC | 143 |
| TM4SF20 | TCCAGGCTCTCTTAAAAGGTCC | ATGGTGTCGTTACTGGTGGG | 181 |
| NR4A3 | GAAGAGGGCAGCCCGGCAAG | ACGCAGGGCATATCTGGAGGGT | 176 |
| PTGS2 | GTTCCCACCCATGTCAAAAC | CCGGTGTTGAGCAGTTTTCT | 108 |
| SIK1 | AGCTTCTGAACCATCCACACA | TTTGCCAGAACTTCTTCCGC | 162 |
| PDE4B | CCGATCGCATTCAGGTCCTTCGC | TGCGGTCTGTCCATTGCCGA | 96 |
| PDE4D | AACACATGAATCTACTGGCTGA | TCACACATGGGGCTTATCTCC | 253 |
| GADPH | CAATGACCCCTTCATTGACC | GACAAGCTTCCCGTTCTCAG | 106 |

**CHIP-PCR primers:**

| Promotor | Forward Primer | Reverse Primer | Amplication (bp) |
|---|---|---|---|
| LINC00473 promotor | CTACAGACGTCATCGCCTCC | CACATTTGGGGGTGCTTGTG | 129 |
| NR4A2 promoter | GGGGAAAGTGAAGTGTCG | CCGCGCTCGCTTTGGTAT | 198 |

**sgRNA target squences:**

| | |
|---|---|
| sgCRTC1-A | TGGCGACTTCGAACAATCCG |
| sgCRTC1-B | TTACCCGCGCGGCCCGCGTC |
| sgCRTC1-C | CCCAGCCGAGGCCAGTACTA |
| sgCRTC2-A | GCAGCGAGATCCTCGAAGAA |
| sgCRTC2-B | AGGATATGTGGCGGGTGTAT |
| sgCRTC2-C | ACAGGCCCAAAAACTGCGAC |
| sgCRTC3-A | CTGACGCACTGCTCCGCAGC |
| sgCRTC3-B | AAAAAGGATATTTGTCGCCC |
| sgCRTC3-C | AACCCGCCATCACGGGCTGG |
| sg-Ctr | CTTCCGCGGCCCGTTCAA |
